# Quantifying Cell Traction Forces at the Individual Fiber Scale in 3D: A Novel Approach Based on Deformable Photopolymerized Fiber Arrays

**DOI:** 10.1101/2024.11.14.623421

**Authors:** Pierre Ucla, Joanne Le Chesnais, Henri Ver Hulst, Xingming Ju, Isabel Calvente, Ludovic Leconte, Jean Salamero, Isabelle Bonnet, Catherine Monnot, Hélène Moreau, Jessem Landoulsi, Vincent Semetey, Sylvie Coscoy

**Author notes:** **Materials and correspondence** should be addressed to P.U., V.S. or S.C.

## Abstract

The forces exerted by cells upon the fibers of the extracellular matrix play a decisive role in cell motility in development and disease. How the local physical properties of the matrix (such as density, stiffness, orientation) affect cellular forces remains poorly understood. Existing approaches to measure cell 3D traction forces within fibrous substrates lack control over the local properties and rely on continuum approaches, not suited for measuring forces at the scale of individual fibers. A novel approach is proposed here to fabricate multilayer arrays of deformable fibers with defined geometrical and mechanical properties using two-photon polymerization. The fibers are characterized using Atomic Force Microscopy and span a wide range of sizes and mechanical properties. This approach is combined with a new reference-free method for measuring traction forces in 3D, which relies on automated segmentation of the fibers coupled with finite element modeling. The force measurement pipeline is applied to study forces exerted by adherent cells, endothelial cells, fibroblasts or macrophages, and reveals how these forces are influenced by fiber density and stiffness. Additionally, coupling to fast volumetric imaging with lattice light-sheet microscopy enables the measurement of the low-intensity and short-lived tractions exerted by amoeboid cells, such as dendritic cells.

## Introduction

Force generation is an essential feature of cell physiology that is important for a variety of processes such as migration, differentiation, and signaling. The mechanical and topographical cues provided by the network of fibrous proteins in the natural extracellular matrix (ECM) regulate cell adhesion, force generation, and, more broadly, mechanotransduction^1^. The relationship between local physical properties—such as matrix density, stiffness, and fiber orientation—and force generation is especially critical since, in pathological conditions like cancer, the ECM is often remodeled, becoming highly heterogeneous^2^. This leads to the generation of 3D gradients of stiffness and fiber density that strongly affect cell fate and behavior. Understanding how local physical properties influence force generation at the scale of individual collagen fibers is fundamental to decipher mechanisms of ECM remodeling, cell migration, and metastasis. However, measuring the forces generated by cells *in situ*^3^ in their native context is challenging and offers little control over the properties of the local microenvironment. In past decades, a large array of tools has been developed to control mechanical properties and measure traction forces on 2D substrates *in vitro*, but it has become increasingly evident that mechanobiology studies should address forces in 3D^4^.

Hydrogels commonly used to study cell mechanobiology in 3D *in vitro* include collagen or fibrin. The traction forces exerted by cells within these hydrogels are usually characterized using traction force microscopy, which measures the displacement of embedded fluorescent beads^5–9^, although it is also possible to measure the displacement field of the fibers themselves using confocal reflectance microscopy^10^.

Despite these developments, important limitations remain in accurately measuring the traction forces generated by cells in 3D hydrogel matrices. First, measurements from cells embedded within collagen- or fibrin-derived hydrogels may lack reproducibility due to the heterogeneous nature of these materials and their degradation by proteases secreted by the cells. Second, with traction force microscopy, the displacement field is calculated by comparing the imaged volume with a stress-free reference state acquired at the end of the experiment; any plastic deformations of the matrix during the experiment can result in an altered reference state and errors in force reconstruction. Third, the hydrogels are typically treated as homogeneous materials and characterized mechanically using large-scale rheology techniques, which does not fully account for the local heterogeneities in properties such as stiffness, pore size or ligand density that cells sense at the scale of individual fibers^11–13^. Engineered hydrogels^14^ made from PEG^15^ or alginate^16,17^ can help fine-tune these parameters, but they have a porous rather than fibrillar architecture. Finally, collagen- or fibrin-derived hydrogels display strong non-linear features due to fiber buckling, straightening or stretching. Material models that consider these features rely on continuum approaches with large representative volume elements which are not suited for measuring forces exerted on individual fibers^7^.

Alternative approaches to produce simplified 3D structures using microfabrication techniques^18,19^ allow for increased control of the local physical properties of the substrate. Yet, their use in cell traction force measurement has largely been limited so far to 2D force measurements. The deflection of arrays of polydimethylsiloxane micropillars produced by stereolithography^20^ can be used for precise force measurement and fine control of the rigidity experienced by cells^21^. Yet, this approach only allows for the measurement of tangential forces and does not account for vertical forces. Alternatively, arrays of polystyrene fibers with submicron diameters, fabricated using the spinneret-based tunable engineered parameters technique, provide a robust framework for fiber micromechanics^22,23^, but this approach is also limited to the measurement of in-plane traction forces^19^.

In contrast to the microfabrication methods described above, two-photon polymerization (TPP) allows tremendous freedom in the geometry, chemistry and mechanical properties of 3D microstructures being fabricated. This technique has been used to produce a variety of 3D microscaffolds in order to study cell migration, morphology and forces in response to architectural cues of the microenvironment^24,25^. However, the range of forces measurable was largely limited by the high structural stiffness of these scaffolds and measurements have been so far restricted to in-plane (2D) forces of cells with high contractility^18,26^.

In this paper, we propose a novel method for fabricating multilayer arrays of highly deformable hydrogel-based fibers with fully controlled geometry and mechanical properties and we combine it with an original pipeline to measure the 3D traction forces locally exerted by cells on these fibers. The range of sizes and mechanical properties of the produced fibers is characterized by Atomic Force Microscopy (AFM). The in-plane and out-of-plane deflections of these fibers submitted to cell traction forces are quantified using fast fluorescence microscopy combined with an automated 3D image analysis framework. The traction forces are then calculated using a regularized inverse method based on finite element analysis. A key advantage of this method is that it does not require a reference to a stress-free state at the end of the experiment. This novel approach provides a versatile tool for life scientists to quantify cell contractility and reconstruct 3D traction force maps of cells confined within a fibrillary microscaffold with reproducible geometrical and mechanical properties. We validate the approach using it to quantify the traction forces of two well-characterized adherent cell types: NIH/3T3 fibroblasts and human umbilical vein endothelial cells (HUVEC), and show how the exerted forces depend on fiber stiffness and density. Furthermore, we demonstrate the utility of our method for studying the low amplitude forces characteristic of immune cells by quantifying the traction forces generated by murine macrophages. Additionally, we show that it can be easily adapted for use with fast volumetric imaging systems, such as lattice light-sheet microscopy, enabling high spatiotemporal resolution in 3D. We apply this approach to quantify the short-lived traction forces exerted by amoeboid cells, such as murine dendritic cells, and highlight the transient anchors and short-lived protrusions that contact and deflect the fibers.

## Results

### Microscaffold design

Two-photon polymerization offers great flexibility for building 3D scaffolds with versatile geometries^27^. We extended the range of this technique to create multilayer arrays of long deformable fibers by developing experimental procedures, including the careful selection of architecture design, materials and processing methods. For simplicity, we focused on a two-layer fiber configuration, with suspended fibers printed orthogonally to maximize the freedom of conformations and spatial exploration of cells subjected to contact guidance on each layer (Fig. 1A-C). Adding extra layers to the microscaffolds posed no technical challenges during the printing process (Supplementary Figure 1). The fiber spacing was tunable in all dimensions, with a lower limit of 5 μm, set by fiber entanglement during fabrication and development. Here, we compared fiber spacings of 10 μm (Fig. 1C-top, D-F) or 5 μm (Fig. 1C-bottom) in the *x-y* plane. In the *z* direction, a spacing of 10 μm was selected, small enough to allow cells to form protrusions between layers, facilitating 3D spreading of cells (Fig.1B). In terms of materials, we designed composite microscaffolds composed of cell-adhesive fibers supported by cell-repellent structural part (walls and carpet) (Fig.1A, B). Resin 1, used for the structural parts, was composed of a mix of Polyethylene Glycol Diacrylate (PEGDA575) with 15% Pentaerythritol tetraacrylate (PETA), making it anti-adhesive, while resin 2, used for the fibers, consisted of PEGDA250 with 10% PETA, allowing protein adsorption^28^ (Supplementary Figure 2). The fibers were then coated with fibronectin coupled with the highly photostable far-red dye CF™ 640R. We seeded NIH/3T3 Lifeact-GFP fibroblasts and HUVEC Lifeact-GFP endothelial cells - two well-characterized adherent cell types - onto these microscaffolds. We confirmed that this 3D scaffold geometry favored cell adhesion, spreading, traction force exertion and migration of these adherent cells on the suspended fibers, with minimal cell adhesion to anti-adhesive structural parts (Fig.1C-F, Supplementary Movies S1,2).

**Figure 1.**
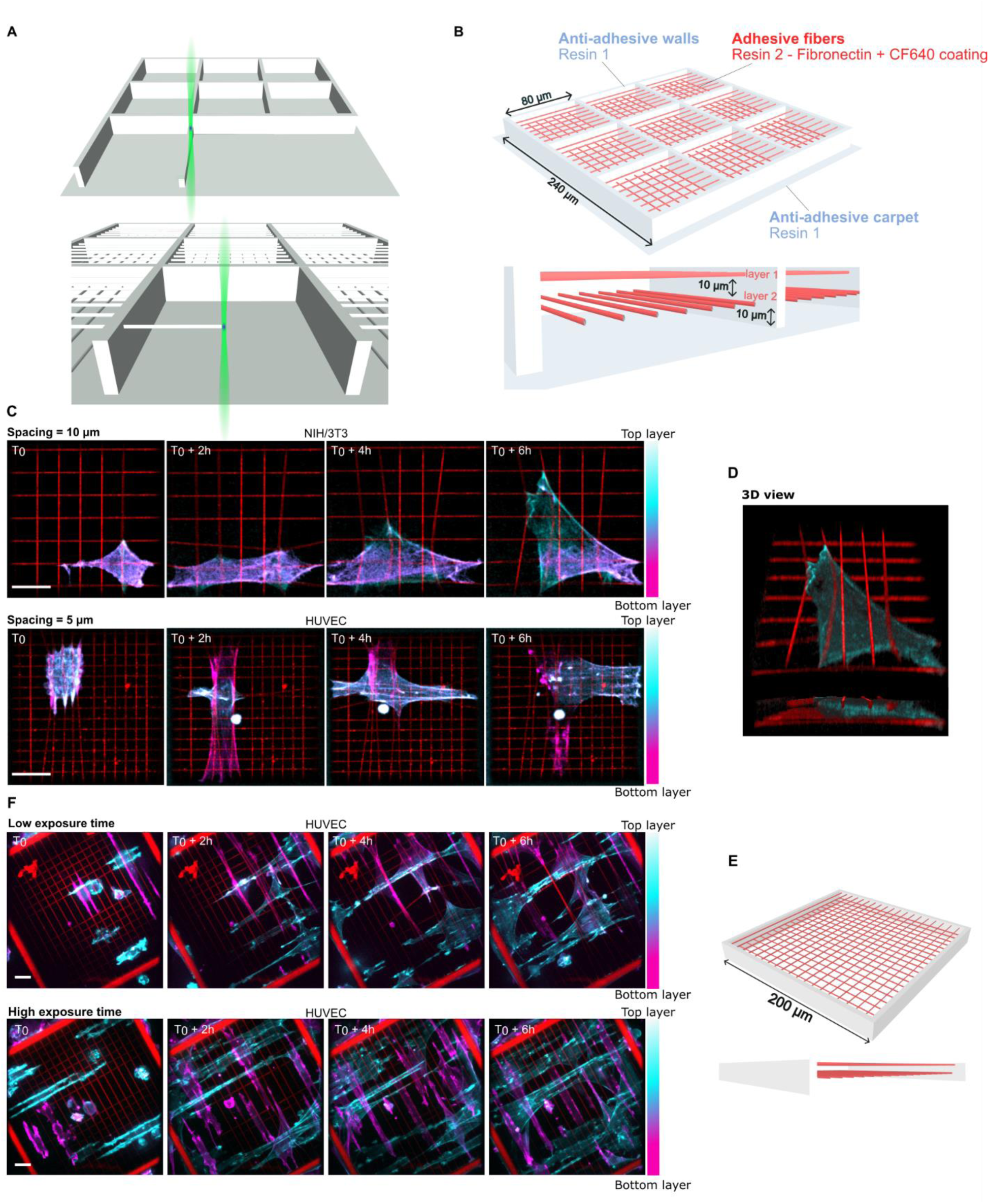
Photopolymerization of multi-layered microscaffolds with fibers. **A.** Two-step photopolymerization process. The structural parts are fabricated first with the anti-adhesive resin (top) before photopolymerization of the fibers as a single voxel line with the adhesive resin (bottom). **B.** Schematic of a 2- layer composite microscaffold with 80 μm fibers after the fabrication and fibronectin coating. Top view (top) and side view (bottom). **C.** Timelapse of a single NIH/3T3 Lifeact-GFP cell spreading and exerting traction forces on fibers with 10 μm lateral spacing (top) and a HUVEC Lifeact-GFP cell on fibers with 5 μm lateral spacing (bottom). (color scale: actin, red: fibers; maximum projection). Scale bar: 20 μm. **D.** 3D reconstruction of an array with NIH/3T3, with top view (top) and side view (bottom). **E.** Schematic of a 2-layer composite microscaffold with 200 μm fibers. **F.** Timelapse of multiple HUVEC Lifeact-GFP cells spreading and exerting traction forces on highly deformable low exposure time photopolymerized fibers (top) and on less deformable high exposure time photopolymerized fibers (bottom). (color scale: actin, red: fibers; maximum projection). Scale bar: 20 μm.

### Geometrical and mechanical characterization of the fibers

A key aspect of our method is the generation of highly deformable fibers with controlled mechanical properties. These properties can be modulated by two fabrication parameters, laser power and scanning speed, with the latter controlling voxel exposure time. We observed that fiber deformability under cellular forces significantly decreased as exposure time increased, as demonstrated by the collective behavior of HUVECs on 200 μm grids (Fig. 1E-F, Supplementary Movies S3,4). This finding suggests that fiber stiffness can be fine-tuned by adjusting voxel exposure time. To build our force reconstruction model, we systematically examined the geometrical and mechanical properties of the photopolymerized fibers across different exposure times.

First, we used Atomic Force Microscopy (AFM) imaging to systematically measure the dimensions of fibers produced at a constant laser power (see Methods), with exposure times ranging from 600 μs to 800 μs (Supplementary Figure 3; see Supplementary Table 1 for the corresponding scanning speeds). Following the procedure described by Buchroithner *et al.*^29^, fibers were collapsed on the substrate prior to AFM imaging. During this process, the fibers rotated along their axis, allowing measurement of the axial and lateral dimensions as the width and height of the collapsed fibers (Fig.2A, B). As expected for photopolymerized structures, the fiber cross-section was elliptical due to diffraction constraints and waveguide effect^30,31^ and the cross-section area increased with increasing exposure times. To quantify the lateral dimension of fibers, we measured both the maximum height and the average height (Fig.2C), the latter being used to approximate a rectangular cross-section in our mechanical model. The maximum height increased from 126 ± 5 nm at 600 μs to 198 nm ± 2 nm at 800 μs and the average height increased from 73 ± 7 nm to 142 ± 4 nm. The axial dimension increased from 1.56 ± 0.03 μm at 600 μs to 2.33 ± 0.04 μm at 800 μs (Fig.2D). The corresponding aspect ratios were relatively high and ranged between 11 and 13, with high aspect ratios already being reported in other studies^29,32^ (Fig.2E). We also observed a periodic nanostructuration on the photopolymerized fibers, with an amplitude and a period decreasing as exposure time increased (Supplementary Figure 4). Similar variations in photopolymerized voxel dimensions have been reported, likely linked to piezo oscillations affecting scanning speed^33^. Next, nanomechanical indentation measurements using force spectroscopy were performed to determine the Young’s modulus of the collapsed fibers (see Methods, Supplementary Figure 5). Some dispersion in the measurements was observed, which we attributed to crosslinking heterogeneity along the fiber profile. The mean value was about 10 MPa (Fig.2F, mean E = 11.32 ± 10.59 MPa, pooling all fiber conditions for a 1 nN load) consistent with previous mechanical characterization of PEGDA and PETA^29,34,35^.

**Figure 2.**
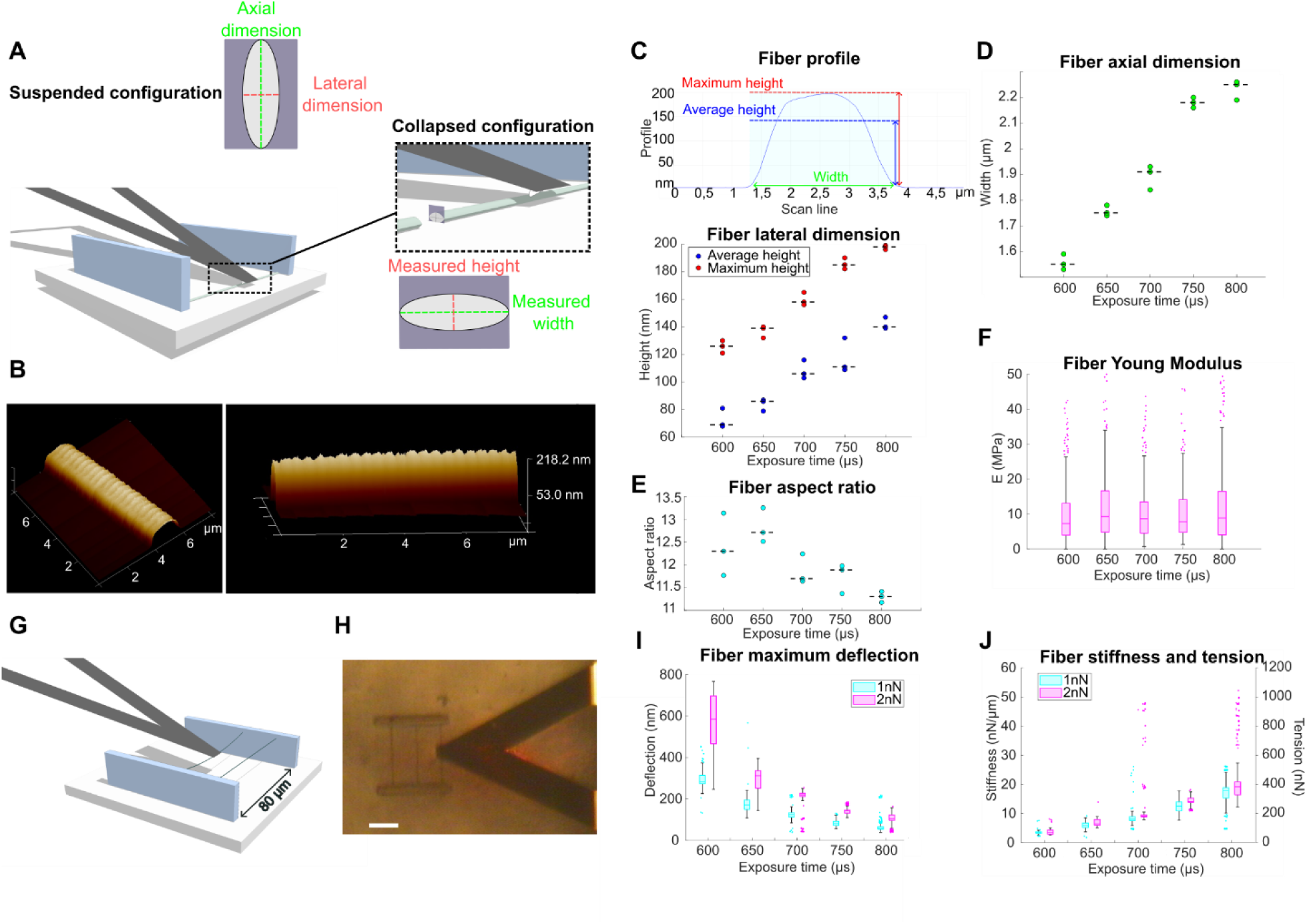
Geometrical and mechanical characterization of the photopolymerized fibers. **A.** Scheme of the AFM topography and indentation measurements, illustrating how the AFM tip is brought in contact with the fiber that is collapsed on the glass coverslip. The fiber has an elliptic cross-section and rotates around its long axis when collapsing. **B.** 3D view of a fiber for T_exp_ = 800 μs (left: top view, right: side view). **C.** (Top) Scanning profile of a fiber for T_exp_ = 800 μs as measured by AFM. The average height corresponds to the mean value of the height profile function over the fiber interval. (bottom) Variation of the lateral dimension of the produced fibers for a range of exposure times (N = 3 fibers for each condition). **D.** Variation of the axial dimension of the produced fibers for a range of exposure times (N = 3 fibers for each condition). **E.** Aspect ratio between axial dimension and maximum lateral dimension of the fibers. For all scatter plots, the dashed line indicates the median. **F.** Young’s Modulus of the produced fibers for a range of exposure times as measured by AFM indentation (N = 3 fibers and n > 280 analyzed force curves for each condition). **G.** Scheme and **H.** bright field image of the suspended fiber deflection experiment, illustrating how the AFM tipless cantilever is brought in contact with the middle point of the suspended fiber (scale bar: 20 μm). **I.** Maximum deflection of the fiber, **J.** corresponding stiffness and tension for an applied force of 1 nN and 2 nN (N = 3 fibers and n = 768 analyzed force-distance curves for each condition). For all box plots, the central mark indicates the median, the edges of the box denote the 25th and 75th percentiles, the whiskers extend to the most extreme data points not considered outliers, and outliers are plotted individually (dots).

For all tested exposure times, we observed shrinkage-induced tension accumulation during the development process with the fibers transitioning from a loose state in the monomer solution to a tense state after washing the uncured monomer with pure ethanol (Supplementary Movie S5). At this stage, fibers can be described as clamped beams under tension. Their apparent stiffness at the midpoint results from the contribution of two components: structural stiffness, which is related to the elasticity of the material and the beam geometry, and stiffness resulting from the tensile loading. When structural elasticity dominates (that is when 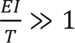), the apparent stiffness is given by 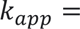 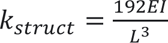. However, when tension dominates (i.e. when 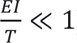), the apparent stiffness becomes 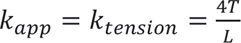 (see Supplementary Note 1). Here, *E* is the Young’s modulus of the material, *I* is the fiber moment of inertia, *L* is the fiber length, and *T* is the tension.

In order to estimate the tension accumulated within the fibers during the development step, we measured the apparent stiffness of suspended fibers by performing force-distance measurements using an AFM tipless cantilever, brought into contact with the midpoint of each fiber (Fig.2G, H). We performed measurements with applied forces of 1 nN and 2 nN and recorded the resulting deflection (Supplementary Figure 6). The deflection decreased as exposure time increased, confirming our ability to tune the fiber stiffness by adjusting fabrication parameters (Fig.2I, J). The corresponding apparent stiffness for a 1 nN force ranged from 3.5 ± 0.4 nN/μm at Texp = 600 μs (3.6 ± 0.7 nN/μm for a 2 nN indentation force, mean ± s.d.) to 17.6 ± 3.6 nN/μm at Texp = 800 μs (respectively 19.8 ± 5.2 nN/μm). This spectrum matches the lower bounds described for pillar arrays^36^ or STEP-deposited fibers^19^ and is suitable for measuring a wide range of cell-generated forces. For clamped fibers, the apparent stiffness measured corresponds to the midpoint of the fiber, with the stiffness profile increasing non-linearly along the fiber length (Supplementary Figure 7). Based on the measured Young’s modulus and fiber dimensions, we estimated that the structural stiffness remained below 0.6 nN/μm for all exposure times studied, accounting for less than 3% of the total apparent stiffness. This indicates that the mechanical regime is largely dominated by tension. Using the average apparent stiffness measured for 1 nN and 2 nN applied forces, we calculated tension values for each condition, which ranged from 71 ± 12 nN for Texp = 600 μs and 374 ± 93 nN for Texp = 800 μs (Fig.2J). The geometrical and mechanical information extracted from the ensemble of AFM measurements was integrated to a pipeline enabling force reconstruction from fiber deflection measurements.

### A constrained reference-free inverse traction recovery framework

We developed a Matlab workflow to compute fiber deflection (Fig.3). Each individual fiber was segmented in 3D based on its fluorescent labelling and tracked over time (see Methods). A deflection vector was calculated as the distance between the deflected fiber skeleton and the straight line connecting its two anchored extremities. This approach offers the advantage being reference-free, eliminating the need for cell trypsination or the use of Cytochalasin D post-experiment to recover the stress-free state. The theoretical resolution of this deflection measurement is limited by the voxel size and could be lowered to the diffraction limit. However, experimental factors – such as imaging system resolution, acquisition frequency, cell sensitivity to phototoxicity or desired computation time – can affect this resolution. Under our experimental conditions, the resolution was 0.3 μm in the *x-y* plane and 1 μm in the *z* direction. These deflection measurements enabled the computation of a dynamic 3D deflection map, showing the local deflection of each fiber within the microscaffold at each time point, and used as the input of an inverse traction force recovery algorithm.

**Figure 3.**
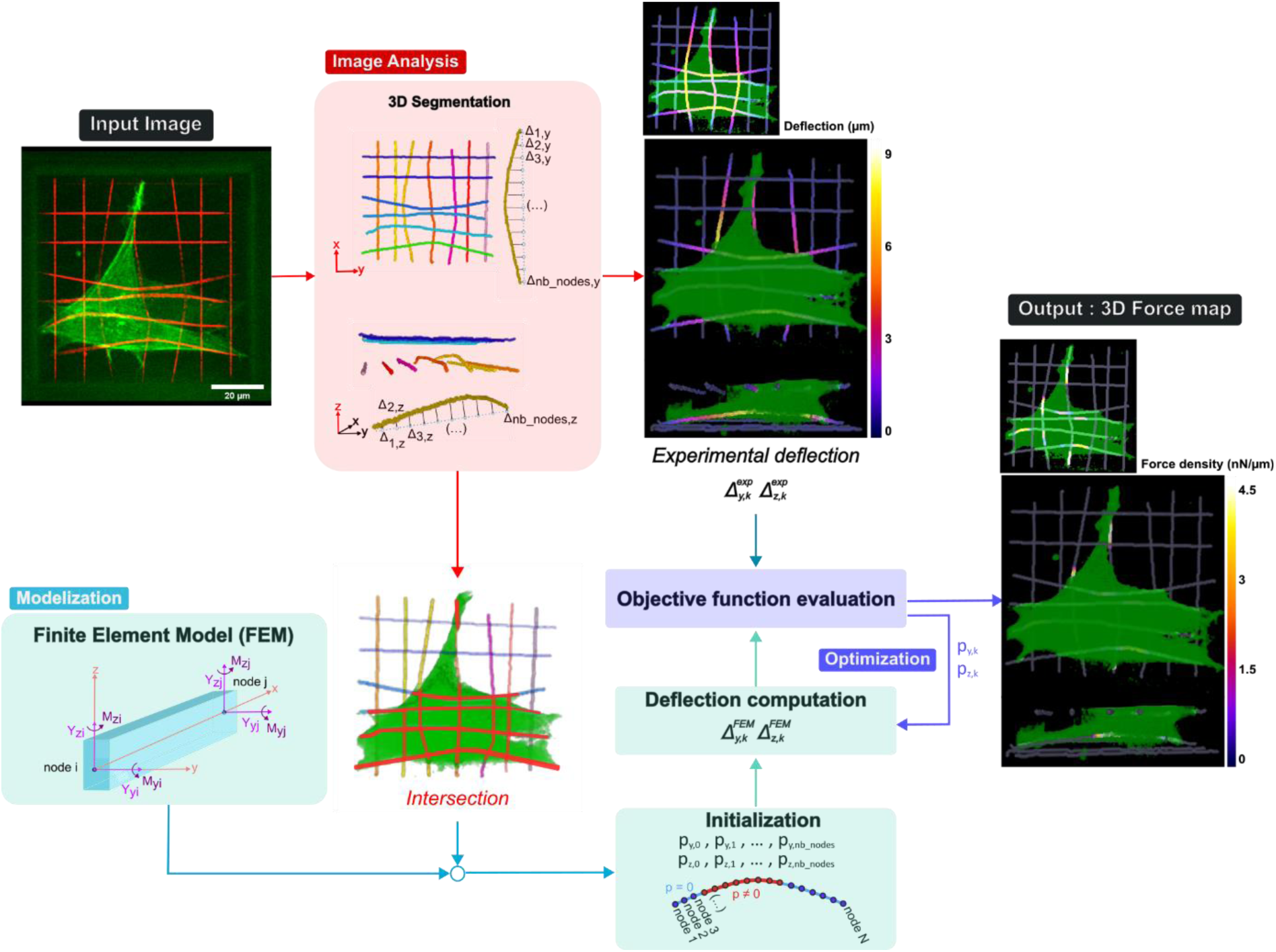
3D Traction forces recovery workflow. The input image stack is first treated by performing the 3D segmentation of the fibers. For each individual fiber, a deflection vector is computed in the two directions orthogonal to the axis of the fiber. The corresponding deflection map is generated where each voxel represents the sum of these two components. The cell volume is also segmented and the intersection between the cell and the fibers is computed to spatially constrain the subsequent traction force recovery. The force vector of a Finite Element Model (FEM) based on 3D beam elements is initialized randomly in the intersection area and an optimization step is performed to minimize the distance between experimental and computed deflections. The output is a force vector for each fiber that is translated into a 3D force map.

Finite Element Modeling (FEM) is traditionally used in 2D and 3D Traction Force Microscopy (TFM) to reconstruct the traction force field exerted on a bulk hydrogel, modeled as a discretized semi-infinite half space. In our case, FEM was used to study the mechanics of individual fibers. Each fiber was treated as a rectangular beam under tensile loading with clamped edges. It was divided into a finite number of 3D beam elements, each having 4 degrees of freedom: two transverse deflections (along y and z axes) and two rotations (around the *y* and *z* axes). Axial extension (in the direction of the fiber axis) and torsion were not considered (see Methods and Supplementary Note 1 for details about the FE model). Ignoring axial forces leads to an underestimation of contractility which we evaluated by looking at the residual forces (Supplementary Note 3). Quantifying this axial contribution could be achieved by incorporating and tracking fluorescent nanoparticles within the fiber resin but it would require acquiring reference undeformed state, thereby eliminating a key advantage of our reference-free method.

The inverse problem of linking fiber deflection to the exerted forces was solved using an optimization framework. The force reconstruction was constrained to the area of intersection between the cell and the fibers. To avoid overfitting the model parameters, we used an elastic net (L1 + L2) regularization approach^37^. The resulting loss function was as follows:

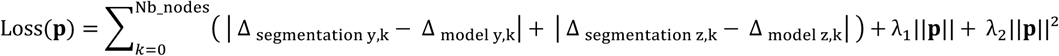

where **p** is the force vector, Δsegmentation y,k and Δsegmentation z,k are respectively the *y* and *z* deflection at node *k* extracted from experimental data, Δmodel y,k and Δmodel y,k are the deflections computed by the model, and λ1 and λ2 are the regularization parameters. The optimal regularization parameters were determined based on simulations, by generating artificial traction force patterns and introducing characteristic measurement noise into the corresponding deflection field (Supplementary Note 2).

### Applications: 3D traction forces maps and cell contractility measurements

Traction forces were first investigated in two classical models of adherent cells known for generating high levels of contractile forces: NIH/3T3 fibroblasts and HUVEC endothelial cells (Fig.4A, B, Supplementary Movies S6-12). These cells spread, occasionally forming very long protrusions around fibers, as previously reported in other fiber-based bioengineered systems^38,39^. Cells contacted both *z* layers with possible transitions between fully spreading across both layers, and predominantly interacting with one layer (Sup. Movie S6-12). Traction force maps showed a spatial distribution of forces, with high force density at the cell periphery and protruding tips and weaker forces in the central region of the cell. Similar patterns of cellular tractions were previously reported on patterned hydrogels^40^, micropost arrays^41^ and suspended fibers^19^ and theoretical models suggested that local traction forces depend linearly on the distance from the cell center^42^. Contractility, measured as the sum of the exerted forces, typically ranged between 50-100 nN for HUVECs and 100-150 nN for NIH/3T3 with a 10 μm lateral spacing (Fig.4C, i and ii). Fig.4C (ii) illustrates the characteristic increase in contractility associated with the spreading of a NIH/3T3 fibroblast over 2-3 hours post-seeding (Sup. Movie S6).

**Figure 4.**
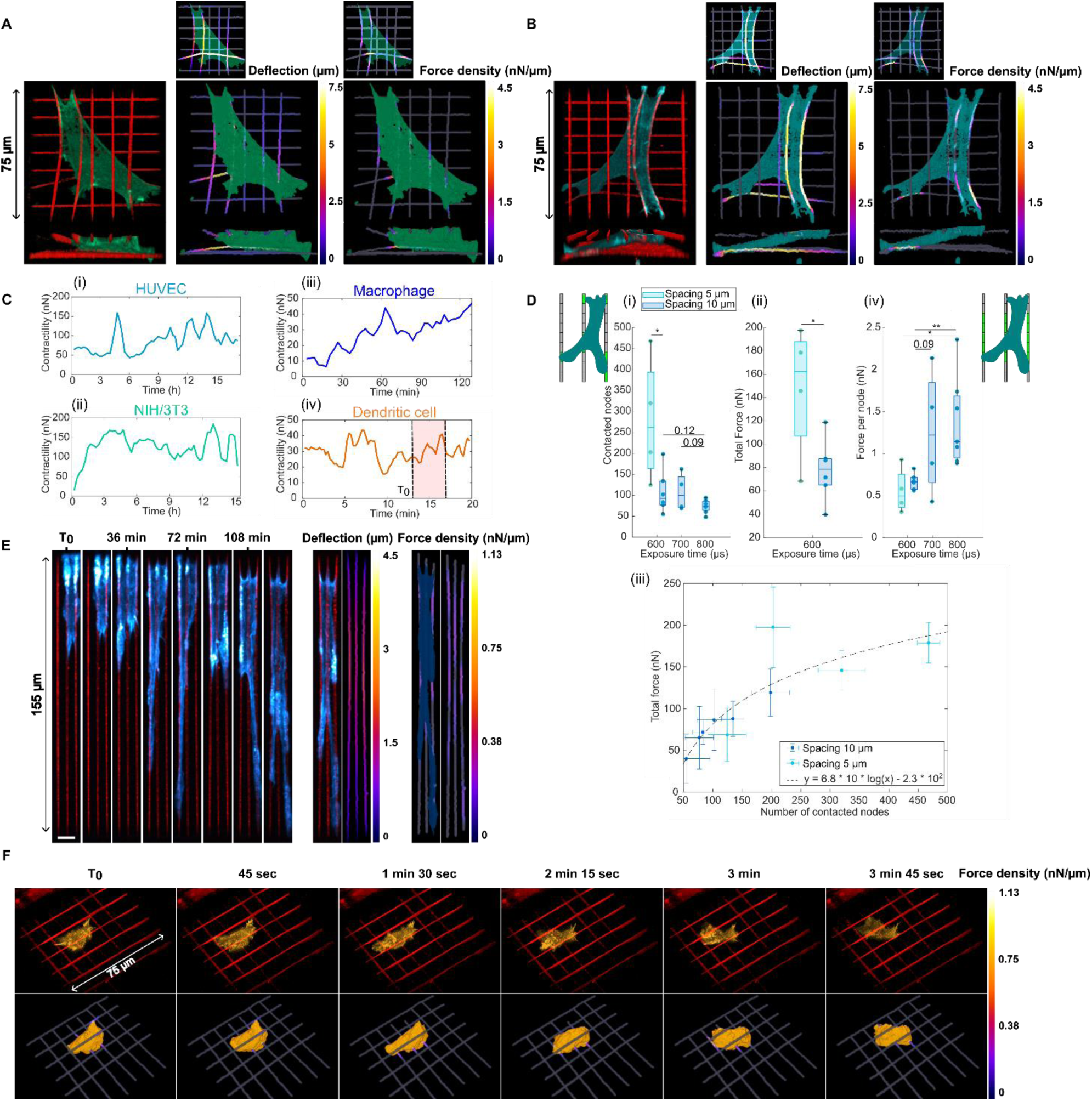
Traction force profiles of various cell lines. 3D reconstruction (left) of a NIH/3T3 cell (**A**) and a HUVEC cell (**B**) on a 2-layer fiber array with 10 μm lateral spacing and low-stiffness fibers (T_exp_ = 600 μs, k = 3.5 nN/μm) with the corresponding 3D deflection maps (center) and 3D traction force maps (right). Side views are represented on the bottom. **C.** Evolution of cell contractility over time for the respective examples of (i) HUVEC shown in **A**, (ii) NIH/3T3 shown in **B**, (iii) macrophage shown in **E** and (iv) dendritic cell shown in **F**. **D.** Effect of fiber exposure time and lateral spacing on (i) average number of contacted nodes, (ii) average total force and (iv) average force per node in the central part of the fiber for HUVECs. Measurements of individual cells are shown as points (600 μs, spacing 5 μm: N = 4; 600 μs, spacing 10 μm: N = 6; 700 μs: N = 4; 800 μs: N = 7). * indicates p < 0.05 and ** indicates p < 0.01 for two-sided t-test. For all box plots, the central mark indicates the median, the edges of the box denote the 25th and 75th percentiles, the whiskers extend to the most extreme data points not considered outliers. (iii) Total force as a function of the number of contacted nodes and logarithmic fit (R² = 0.76) for HUVECs on low-stiffness fibers (T_exp_ = 600 μs, k = 3.5 nN/μm) with lateral spacings of 5 μm (cyan) and 10 μm (blue). Each point represents the mean value over the whole timelapse for an individual cell and error bars correspond to the standard deviation. **E.** Timelapse illustrating adhesion and spreading of a macrophage Lifeact-GFP on one layer of medium-stiffness fibers (T_exp_ = 700 μs, k = 4.1 nN/μm) with lateral spacing of 5 μm while deflecting the fibers (left) (blue: actin, red: fibers). Scale bar: 10 μm. Deflection map for one timepoint (center) and corresponding force map with segmented cell mask (right). **F.** Lattice light-sheet timelapse of a dendritic cell migrating on a 2-layer fiber array with 10 μm lateral spacing and low-stiffness fibers (T_exp_ = 600 μs, k = 3.5 nN/μm) (top) (orange: actin, red: fibers) and corresponding 3D traction force map (bottom).

We sought to evaluate the ability of our system to modulate cell mechanics based on the local topography and mechanical properties of the fiber arrays. Specifically, we examined the influence of fiber density and stiffness on adhesion and traction forces generated by HUVECs. Cells were seeded on low-, middle- or high-stiffness fibers with a lateral spacing of either 5 or 10 μm. We quantified cell adhesion by measuring the number of contacted nodes on the discretized fibers. On fibers with 5 μm lateral spacing, cells contacted more than twice the number of nodes (average of 279) compared with 10 μm fiber spacing (average of 108) (Fig.4D, i). For low-stiffness fibers, we observed a significant increase in total exerted force with fiber density, from 78 ± 26 nN for 10 μm spacing to 147 ± 57 nN for 5 μm spacing (Fig.4D, ii). This was associated with a logarithmic relationship between the total exerted force and the number of contacted nodes (Fig.4D, iii). Accordingly, the average force per node decreased with increasing number of contacted nodes (Supplementary Figure 8), in agreement with previous studies on the effect of micropost density^41^. Lastly, we examined the influence of fiber stiffness on the generated forces. We observed that the average force per node in the central area of the fiber (Supplementary Figure 7) increased with fiber stiffness from 0.67 ± 0.10 nN/node to 1.4 ± 0.54 nN/node, respectively for low- and high-stiffness fibers with a 10 μm spacing (Fig.4D, iv). This suggests that similar force-stiffness relationships govern traction forces in both 3D fibrillary systems and in simpler 2D systems^19,43,44^. It is important to note, however, that the increase in fiber cross-section area with exposure time could also contribute to this increase of local forces. Altogether, these results demonstrate that both fiber density and stiffness modulate cell traction forces, influencing not only local forces but also the larger-scale mechanics across the entire fiber array.

To assess the potential of our method for measuring the lower range of traction forces typically generated by immune cells, we seeded macrophages Lifeact-GFP on very low-stiffness fibers, reducing their apparent stiffness by doubling the fiber length from 80 μm to 160 μm. Macrophages predominantly remained on the upper layer of the scaffolds. They adopted a highly elongated morphology with dynamic protrusions and contacted only two to three fibers at most, for the two lateral spacing considered (5 and 10 μm) (Fig.4E) (Supplementary Movies S13-16). Measurable deflections in the range of 1-2 μm were observed, with measured contractility values between 10-50 nN (Fig.4C, iii), consistent with reported traction forces for leukocytes^9^. Finally, 3D force measurements of extremely dynamic cell types remain challenging and require good compatibility with ultrafast volumetric imaging methods. To demonstrate the suitability of our setup, we combined it with Lattice Light-Sheet imaging to study the migration and force generation of dendritic cells on low-stiffness fibers. Although the motility of dendritic cell is classically described as amoeboid and adhesion independent^45,46^, we observed transient anchoring of the cells on the fiber *via* their uropod, and the formation of short-lived protrusions contacting and deflecting the fibers (Supplementary Movies S17,18) (Fig.4F). In contrast to macrophages, dendritic cells maintained a more rounded morphology and could frequently contact both layers of the scaffold. We measured a contractility in the range of 10-40 nN, consistent with previous studies on micropost arrays^47,48^, although slightly higher than what was measured on other suspended fiber systems^49^, likely due to higher fiber density of our multilayer system. The compact shape of DCs resulted in higher local force density than for macrophages, and they also exhibited quick contractility variations, in the range of 1-5 minutes, similar to what was observed for natural killer cells in reconstituted collagen matrices^50^.

## Discussion

In this study, we present a novel technique for fabricating multilayer arrays of highly deformable fibers tailored for traction force measurements. The first innovation of our approach is the development of experimental procedures to obtain multilayer arrays of highly deformable fibers using two-photon polymerization (TPP). The low stiffness of the produced fibers greatly extends the range of forces measurable compared to previous TPP-based methods. The second innovation lies in combining this fiber arrays with an integrated approach for measuring local 3D forces exerted by cells within the scaffolds, providing precise spatial localization of forces and enabling automation of the method. The precise control over 3D geometry and local chemistry afforded by this method enabled us to confine individual cells within simple, defined arrays of microfibers to measure their adhesion and force generation in a 3D fibrillar environment. We demonstrate that the stiffness of the fabricated fibers can be finely tuned across a broad range of values by adjusting polymerization parameters, making them well-suited for studying the deformations induced by various cell types. By developing image analysis tools along with mechanical modeling and a force inference pipeline, we generated 3D traction force maps at the scale of individual fibers, and we computed dynamic measurements of cell contractility. We validated our approach by applying it to two established models of adherent cells with high contractility, namely HUVECs and NIH/3T3 cells. Importantly, we demonstrated how both fiber density and stiffness impact HUVEC adhesion and exerted forces in our 3D fiber system. In addition, our method could be used in conjunction with lattice light-sheet microscopy to measure weaker and highly dynamic forces exerted by immune cells such as macrophages and dendritic cells, and we showed that these latter amoeboid cells form transient anchors and short-lived protrusions that contact and deflect the fibers.

An important point to consider is how the dimensions and nanotexturation of the photopolymerized fibers compare to natural matrices. Collagen fibers *in vivo*, for example, have a wide range of diameters, ranging from several tenths of nanometers for single fibrils^51^ to several microns for bundled fibers^52^. In our set-up, chemical and polymerization parameters control the resolution, anisotropy, and nano-structuration of the fibers produced. Fibers reported in this work have adjustable dimensions of the same order as those of the extracellular matrix. However, we observed an asymmetry in the fiber cross-section, inherent to the anisotropic nature of the two-photon polymerization process. Although some works utilizing two-photon polymerization have reported features in the *z*-axis smaller than 500 nm^29,53^, ensuring stable ultra-thin long fiber arrays at this scale remains a challenge that would require further design on resin chemistry, mechanics, and scaffold architecture. Furthermore, periodic nanostructuring similar to the D-band periodicity^54^ observed in physiological collagen I fibrils - typically around 67 nm - was seen in our fibers by AFM. In our case, the nanostructuration depends on the photopolymerization scanning speed and could likely be modulated to produce either smaller periods similar to those of collagen fibers, or very smooth fibers like those that are typically produced with bioengineering techniques like electrospinning. This flexibility could allow to further study the effect of local nanotexturation on cell response, with potential implications on curvature sensing^55^, adhesion formation^56^ and protrusion dynamics^57^.

Beyond the properties of individual fibers, a fundamental advantage of our approach is its ability to modulate collective fiber organization in 3D. In the current study, we used two perpendicular layers of aligned fibers to create a simple scaffold for measuring 3D forces. Future developments could involve engineering more intricate and physiologically relevant architectures, such as crosslinked fiber networks with diverse 3D fiber orientations. Extending our 3D force measurement method to these complex networks could be highly valuable for validating predictions from computer models on cell migration and force generation within discrete fiber networks^58^. Indeed, control over the key local parameters of fiber networks such as connectivity, length and orientation of the fibers cannot be achieved with standard reconstituted hydrogels. Additionally, the versatility of two-photon polymerization chemistry offers further development opportunities. While our current system uses purely elastic PEGDA fibers, our method could be expanded to incorporate viscoelasticity, which is central in morphogenesis or oncogenesis^59^. This might be achieved by chemically adjusting crosslink strength or, alternatively, by photopolymerizing natural polymers such as ECM proteins, which exhibit viscoelastic behavior. Finally, the design of these networks could also include 3D gradients of local stiffness or fiber density, in an effort to mimic the heterogeneous physical properties found in physiological and pathological ECM. Such designs hold significant potential for creating synthetic 3D tumor microenvironment models to study the effect of local physical cues on cancer, immune and stromal cells.

## Materials and methods

### Photopolymerization setup

The two-photon polymerization set-up consisted of a QSwitch Teem Photonics laser (Grenoble, France), 10 kHz, 5 ns pulses, 10 µJ, 532 nm, a IX70 microscope with a water objective 60× (NA 1.2) LPlanApo, Olympus, and an oil objective 100× (NA 1.4), a piezo-*z* stage and a 3D stage (Physik Instrumente, Karlsruhe, Germany), and a Guppy CCD camera for monitoring structure formation. It was driven by Lithos software, with an autofocus module.

### Microscaffolds preparation

The cell-repellent Resin 1 was made of a mixture of Polyethylene glycol diacrylate (PEGDA, Mn = 575 g/mol, Sigma Aldrich) with 15% (w/w) Pentaerythritol tetraacrylate (PETA, Sigma Aldrich) and 5% (w/v) 2,2-Dimethoxy-2-phenylacetophenone (Irgacure 651, Sigma Aldrich). The cell-adhesive Resin 2 was made of a mixture of Polyethylene glycol diacrylate (PEGDA, Mn = 250g/mol, Sigma Aldrich) with 10% (w/w) PETA and 5% (w/v) Irgacure 651. We checked for selective cell adhesion using simple 2D composite structures (Supplementary Figure 1).

Adhesion of acrylic-based photoresists to the glass substrate surface was enhanced by functionalizing the 30 mm glass coverslips with 3-(trimethoxysilyl)propyl methacrylate (Sigma Aldrich; 1 mM in ethanol) for 5 minutes. Walls of the microscaffold were fabricated with Resin 1 using a 60× water objective (NA 1.2) using a power of 6.4 mW. The excess Resin 1 was removed by washing the coverslip with pure ethanol and air-dried for 1 minute. Resin 2 was then dropcast on top of the fabricated walls. Repositioning of the sample was handled by fabricating two landmarks with Resin 1 during the first step. Fibers were built using a 100× oil objective (NA 1.4) and a power of 4.9 mW. The coverslip was then placed in an incubation chamber and immersed in ethanol to remove the excess Resin 2 and sterilize the sample. This configuration allows the sample to be kept in liquid conditions at all times to prevent collapse of the fibers due to surface tension effects. The sample was then immersed in Phosphate Buffer Solution (PBS) by performing 10 successive partial washings of half the volume of the incubation chamber. The sample was then incubated in fibronectin (Sigma Aldrich, 10 µg/mL) coupled with CF™ 640R succinimidyl ester (Sigma Aldrich) at 37°C for 1 hour. Finally, the sample was immersed in cell medium by performing 10 successive partial washings.

### Cell culture

NIH/3T3 cells, ATCC CRL-1658, were transfected by PEI MAX (Polysciences) with Lifeact-GFP plasmid (pLVX LifeAct-eGFP-P2A-puro, constructed from pLVX puro vector Clontech Catalog No. 632164), selected with 1 µg/ml puromycin and sorted by FACS. NIH/3T3 Lifeact-GFP cells were maintained in DMEM + 10% calf serum (ATCC 61965-026) + 1 µg/ml puromycin, at 37°C, 5% CO2. HUVECs were prepared from human umbilical cords provided by AP-HP, Hôpital Saint-Louis, Unité de Thérapie Cellulaire, CRB-Banque de Sang de Cordon, Paris, France. HUVEC Lifeact-GFP cells were obtained from these cells by transduction with rLVUbi–LifeAct®–TagGFP2 (Ibidi, Gräfelfing, Germany). After thawing, cells were maintained for 1–2 passages in endothelial cell growth medium 2 (ECGM2, Promocell GmbH, Heidelberg, Germany) with 1% P-S, at 37 °C and 5% CO2, in collagen-coated flasks. Dendritic cells were obtained as previously described^60^ by differentiating bone-marrow cells from LifeAct-GFP mice during 10-11 days in IMDM medium (Sigma-Aldrich, Darmstadt, Germany) containing 10% FBS decomplemented and filtered (Biowest, Nuaillé, France), 20 mM L-glutamine (Gibco, Waltham, Massachusetts, USA), 100 U/ml penicillin–streptomycin (Gibco, Waltham, Massachusetts, USA), 50 μM 2-mercaptoethanol (Gibco, Waltham, Massachusetts, USA), and 50 ng/ml of GM-CSF containing supernatant obtained from transfected J558 cells^61^. Only semi adherent (dendritic cells) were recovered at the end of the culture (excluding non-adherent granulocytes and adherent macrophages). Macrophages were obtained by differentiating bone marrow cells from LifeAct-GFP mice in RMPI-Glutamax (Gibco) containing 10% FBS, 1% Pen/Strep, 10 mM Hepes, 1mM NaPyruvate, 50uM B-mercaptoethanol, and supplemented with 30% of M-CSF-containing L929 cell supernatant. Cells were let to differentiate for 7 days before removing M-CSF, and were used on day 8 or 9 after detachment with Trypsin.

### Imaging – Spinning disk

Live imaging experiments were performed using a spinning disk from the PICT-IBiSA platform, equipped with an inverted Eclipse Ti-E (Nikon, Tokyo, Japan) and a spinning disk CSU-X1 (Yokogawa) integrated in Metamorph software by Gataca Systems, with a Prime BSI camera (Photometrics, Tucson, AL, USA) with 6.5 μm pixel size and a z-motor Nanoz100 (Mad City Lab, Madison, WI, USA), temperature and CO2 controllers from Life Imaging Systems, and with a Plan fluor 40X oil (NA 1.3) objective. Image stack size was 180×280×21 μm with a voxel size of 0.16 x 0.16 x 1 μm. For imaging HUVECs and NIH/3T3 cells, a time step of ΔT = 15 min was employed for a total imaging time of 15-17 hours, whereas for macrophages, it was set to ΔT = 3 min and the total imaging time was 2 hours.

### Lattice Light Sheet imaging

Image acquisition was performed on a commercialized version of a previously described setup from 3i (Denver, USA). Cells were scanned incrementally through a 20 μm long light sheet, in 1000 nm steps using a fast piezoelectric flexure stage (equivalent to ≃ 650 nm voxel size in z axis after image realignment, respecting the 32,8° angle for the detection objective position). Data were imaged using sCMOS camera (Orca-Flash 4.0; Hamamatsu, Bridgewater, NJ). Excitation was achieved with 561 nm or 642 nm (MPB Communications, Montreal, Canada) diode lasers at ∼5% acousto-optic tunable filter transmittance, with 50 mW nominal power. Excitation is done through a water-dipping 28.6× objective (NA=0.7, working distance 3.74-mm) and detection *via* a Nikon CFI Apo LWD 25× water-dipping objective (NA=1.1), completed with a 2.5× tube lens, to obtain a final pixel size of 104 x 104 nm. Lattice light-sheet imaging was performed using an excitation pattern of outer NA equal to 0.55 and inner NA equal to 0.493. Composite volumetric datasets were obtained using ≃ 10 ms exposure/optical planes/channel at a time resolution of 15 seconds per total cell volume of 80 planes with a field size of 1200×1200 pixels. Double labeled images were acquired on two separated cameras, using a dichroic mirror with an edge at 660 nm. 3D+time series were acquired within 20 minutes. Acquired data were deskewed, a necessary step to realigned the image frames, using LLSpy, a python library (copyright to T. Lambert, Harvard Medical School, Boston, USA; https://github.com/tlambert03/LLSpy) and deconvolved using cudaDeconv (copyright to Lin Shao et al, HHMI https://github.com/dmilkie/cudaDecon), included in LLSpy. For Lifeact-GFP cells on a two-layer microscaffold, LLSM images were denoised using ND-SAFIR software (https://gitlab.inria.fr/serpico/ndsafir_bin) before deskewing and deconvolution, as above. Napari, a multi-dimensional image viewer for Python (https://github.com/napari/napari).

### AFM experiments

All AFM experiments were performed using a commercial AFM (NanoScope VIII MultiMode AFM, Bruker Nano Inc-Nano Surfaces Division, Santa Barbara, CA). Glass substrates were fixed on a steel sample puck using double-sided tape. All experiments were performed at room temperature (∼22°C) in a Phosphate Buffer Solution (PBS) adjusted at pH 7.4. The cantilevers and samples were left to equilibrate for about 30 min in PBS before starting the measurements.

#### Imaging and nanoindentation

Prior to imaging, long fibers (L = 300 μm) were photopolymerized near the glass substrate (3 μm above the coverslip) and collapsed during the washing steps. The sample was then immersed in PBS and all the subsequent steps were performed in PBS. AFM images were recorded using ozonated oxide-sharpened microfabricated Si3N4 AFM tip purchased from Bruker (SCANASYST-AIR, Nano Inc., Nano Surfaces Division, Santa Barbara, CA). For nanoindentation measurements, the spring constant of the cantilever was measured using the thermal noise method, yielding a value of 0.69 N/m. The curvature radius of silicon nitride tips was about 2 nm (manufacturer specifications). Series of measurements were performed with a constant indentation force of 1 nN. 16×16 windows were used with a scan size of 2 to 5 μm centered around the fiber. The Nanoscope Analysis software (Bruker Nano Inc-Nano Surfaces Division, Santa Barbara, CA) was used to automatically extract the contact point. Poisson ratio was set to 0.5 and the Sneddon model was used to determine the elastic modulus. The applied force was low enough to keep the indentation small and avoid probing the underlying glass substrate (Supplementary Figure 4).

#### Fiber deflection

For mechanical measurements on suspended fibers, a simplified version of the scaffold was designed with only three fibers, suspended 10 µm above the glass substrate. The samples were kept immersed in PBS at all times to avoid fibers collapsing due to surface tension effects. The mechanical measurements were performed using AFM tipless cantilevers, purchased from Bruker (NP-O10, Nano Inc., Nano Surfaces Division, Santa Barbara, CA). The spring constants of the cantilevers were measured using the thermal noise method, yielding values ranging between 0.1297 and 0.1785 N/m. Series of measurements were performed using a 16×16 window without a lateral displacement of the cantilever, on the center of the suspended fibers, using a maximum force of 1 nN or 2 nN. The Nanoscope Analysis software (Buker Inc) was used to automatically extract the contact point using a linear fit.

### 3D segmentation and tracking of the fibers and deflection measurement

#### Pretreatment

Pretreatment of the acquired 3D timelapses was performed using the ImageJ software and consisted of drift correction using the 3D Drift Correction plug-in, resampling with bilinear interpolation to achieve a final isotropic voxel size of 0.3 μm and background subtraction. 3D segmentation, tracking, deflection calculation and force computation were carried out using a custom Matlab script.

#### 3D segmentation and tracking

Briefly, the volumetric image was thresholded and the extremities of fibers were segmented in a semi-automated manner for the first time point. For each subsequent time point, the same threshold was applied to the image, and fibers were segmented as individual connected components. A lower volume threshold was applied to filter small components, while an upper volume threshold was also set to exclude contacting fibers, which could occur in cases of strong deflections. These cases were treated separately, and the two connected fibers were split by computing the minimum geodesic distance between the two extremities of each fiber. Fiber tracking was performed by matching objects with the higher overlap between time points.

#### Deflection measurement

For each time point and each fiber, the local deflection amplitude was computed as the Euclidean distance between the skeletonized deflected fiber and the straight line connecting its two extremities of the fiber. This distance was evaluated separately along the y and z directions for each voxel, and the resulting vector was then subsampled to match the number of nodes used in the FEM model.

### Finite Element Modeling

Each individual fiber was modeled as a rectangular beam under extensive stress and discretized into 3D beam elements, with N nodes of size *l*. The number of nodes was chosen as a trade-off between accuracy of the force reconstruction and computation time and was fixed at 41 (Supplementary Note 2). For experiments involving HUVECs, NIH-3T3 and Dendritic Cells, the node size was 2 μm while for Macrophages, it was 4 μm. Four degrees of freedom were considered: two transverse deflections, *u*_*y*_ and *u_z_*, and two rotations, θ_*y*_ and θ_*z*_, around the y and z axes. The total energy *U*_*tot*_ was calculated as the sum of the strain energy *U_e_* and the potential energy related to the extensive stress *U_e_*: *U*_*n*_ = *U_tot_* + *U_n_*. Using Castigliano’s theorem, the stiffness matrix [*K*] that links the applied stress {*F*} to the resulting strain {δ} was decomposed into [*K_e_*] and [*K_n_*] :

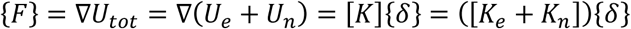

The resulting stiffness matrices were:

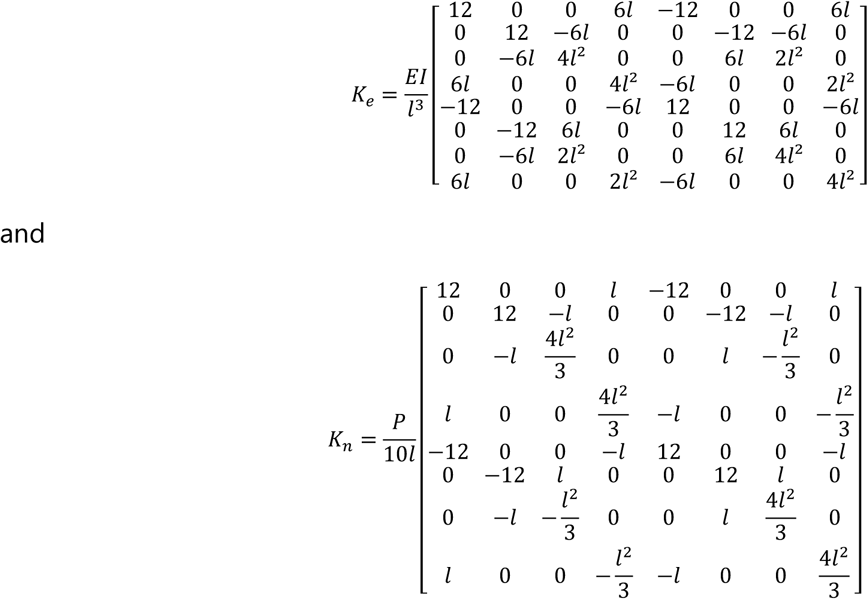

The detailed calculation of the stiffness matrices can be found in Supplementary Note 1.

Computational implementation of the FE model was performed by adapting the Matlab toolbox CALFEM (https://github.com/CALFEM/calfem-matlab). Specifically, the stiffness matrix provided in the “beam3e” function was replaced by the stiffness matrix described above.

## Supporting information

Supporting Movies, Part 1, S1 to S5

Supporting Movie, Part 2, S6 to S11

Supporting Movie, Part 3, S12 to S18

## Code availability

The Matlab code developed for the analyses performed in this paper is available on GitHub : https://github.com/uclapierre/CellFLEX-FM.

## Acknowledgements

The authors acknowledge Axel Buguin, Pascal Silberzan and all the members of the PBME group for their support, helpful discussions and critical reading of the manuscript. We are grateful to François Amblard, Pascal Martin, Carles Blanch Mercader, Stéphanie Descroix, Ana-Maria Lennon-Dumenil, Claire Wilhelm, Patricia Bassereau, Nastassia Pricoupenko, Matthieu Piel and Stéphanie Lebreton for insightful discussions and to Elnaz Nematollahi, Monica Lim, Fanny Tabarin and Fan Sun for their participation in the experiments. We thank Laurent Muller for providing us the HUVEC-Lifeact-GFP cells. We acknowledge Plume Scientific Communication Services (Carol Featherstone) for editing assistance during the preparation of the manuscript. The authors greatly acknowledge the Cell and Tissue Imaging core facility (PICT-IBiSA), Institut Curie, member of the French National Research Infrastructure France-BioImaging (ANR10-INBS-04), and Chloé Guedj for help with imaging experiments.

## Author Contributions

P.U., I.B., C.M., H.M., J.L., V.S., and S.C. designed the study; P.U., J.L.C., I.C., X.J. and L.L. performed experiments; P.U., H.V.H, J.L.C. and L.L. analyzed data; P.U. and S.C. drafted and revised the manuscript; all authors approved the final version of the manuscript.

