## Supplementary Notes for "Quantifying Cell Traction Forces at the Individual Fiber Scale in 3D: A Novel Approach Based on Deformable Photopolymerized Fiber Arrays"

### Supplementary Notes, Figures and Table

#### Supplementary Note 1 - Bending of a beam under extensive stress : analytical solution and FEM formulation

##### I. BENDING OF A BEAM UNDER EXTENSIVE STRESS

In this section, we study the response of a beam that is held under an extensive stress $T$ at both ends to a point load $P$ applied at a position $X = C$ along the beam longitudinal axis $X$, as represented on Sup.Fig.N1.1. To do so, we follow the steps that are detailed in paragraph 4 of Ref. [1]. The load induces a deformation of the beam $Y(X)$ perpendicular to the beam axis. At $X = 0$ (resp. $X = L$), the beam is subjected to an external moment $M_0$ (resp. $M_1$). The equation for the balance of the moments is

$$EI\frac{d^2Y}{dX^2} = TY - P\left(1 - \frac{C}{L}\right)X + M_0 \qquad \text{if } X < C, \tag{S1}$$

$$EI\frac{d^2Y}{dX^2} = TY - P\left(1 - \frac{X}{L}\right)C + M_1 \qquad \text{if } X > C, \tag{S2}$$

$$Y(0) = Y(L) = 0, \tag{S3}$$

$$\lim_{X\to C^-} Y(X) = \lim_{X\to C^+} Y(x), \tag{S4}$$

$$\lim_{X\to C^-} \frac{dY}{dX}(X) = \lim_{X\to C^+} \frac{dY}{dX}(X), \tag{S5}$$

where we introduced the Young modulus $E$ and the moment of inertia $I$ of the beam. In the following, we use the distance $L$ between the ends of the beam as the length scale, and the extensive stress $T$ exerted on both ends of the beam as the stress scale. We define the dimensionless variables $x = X/L$, $y = Y/L$. The problem has three dimensionless parameters: $\alpha = \frac{L}{\lambda}$, where $\lambda = \sqrt{EI/T}$ is a characteristic length, $\beta = \frac{P}{T}$, and $c = C/L$. Finally, we define the two integration constants $m_0 = \frac{M_0}{TL}$ and $m_1 = \frac{M_1}{TL}$, which correspond to dimensionless moments. The resulting set of equations is

$$\frac{1}{\alpha^2}\frac{d^2y}{dx^2} - y = -\beta\,(1-c)\,x + m_0 \qquad \text{if } x < c, \tag{S6}$$

$$\frac{1}{\alpha^2}\frac{d^2y}{dx^2} - y = -\beta\,(1-x)\,c + m_1 \qquad \text{if } x > c, \tag{S7}$$

$$y(0) = y(1) = 0, \tag{S8}$$

$$\lim_{x\to c^-} y(x) = \lim_{x\to c^+} y(x), \tag{S9}$$

$$\lim_{x\to c^-} \frac{dy}{dx}(x) = \lim_{x\to c^+} \frac{dy}{dx}(x). \tag{S10}$$

The sum of the homogeneous and particular solution of Eqs. (S6) and (S7) is

$$y(x) = \begin{cases} C_1\cosh(\alpha x) + C_2\sinh(\alpha x) + \beta\,(1-c)\,x - m_0 & \text{if } x < c \\ C_3\cosh(\alpha(1-x)) + C_4\sinh(\alpha(1-x)) + \beta\,(1-x)\,c - m_1 & \text{if } x > c, \end{cases} . \tag{S11}$$

where the constant $C_i$ must be determined by the boundary conditions. From $y(0) = 0$ and $y(1) = 0$, we respectively find $C_1 = m_0$ and $C_3 = m_1$. The continuity of $y(x)$ and its derivative $\frac{dy}{dx}(x)$ at $x = c$

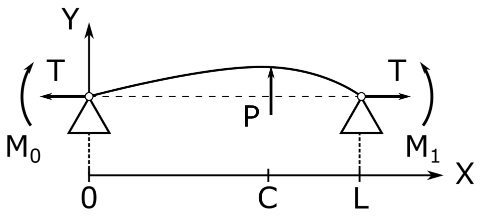

Sup.Fig.N1.1: Sketch of a beam subjected to a point load $P$ applied at the position $X = C$. At its edges, the beam is subjected to an extensive stress $T$ and a moment $M_0$ at $X = 0$ and $M_1$ at $X = L$.

allows to compute the constants $C_2$ and $C_4$. For simplicity, we use the linearity of the problem and write the solution as the sum of the solutions for $\beta$, $m_0$, and $m_1$ treated separately. Let us first consider the case $m_0 = m_1 = 0$. In this setting, the solution of the equation is

$$y_\beta(x) = \begin{cases} \beta\left[(1-c)\,x + C_{\beta,<}\sinh(\alpha x)\right] & \text{if } x < c \\ \beta\left[(1-x)\,c + C_{\beta,>}\sinh(\alpha(1-x))\right] & \text{if } x > c \end{cases}. \tag{S12}$$

with

$$C_{\beta,<} = -\frac{1}{\alpha(\cosh(\alpha c) + \coth(\alpha(1-c))\sinh(\alpha c))}, \tag{S13}$$

$$C_{\beta,>} = -\frac{1}{\alpha(\cosh(\alpha(1-c)) + \coth(\alpha c)\sinh(\alpha(1-c)))}. \tag{S14}$$

Let us now consider the case where $\beta = 0$ and $m_1 = 0$. In this setting, the solution of Eqs. (S6) to (S10) is

$$y_{m0}(x) = \begin{cases} m_0\left[\cosh(\alpha x) - 1 + C_{m0,<}\sinh(\alpha x)\right] & \text{if } x < c \\ m_0 C_{m0,>}\sinh(\alpha(1-x)) & \text{if } x > c \end{cases}. \tag{S15}$$

with

$$C_{m0,<} = \frac{\cosh(\alpha(1-c))}{\sinh(\alpha)} - \coth(\alpha), \tag{S16}$$

$$C_{m0,>} = \frac{1-\cosh(\alpha c)}{\sinh(\alpha)}. \tag{S17}$$

Conversely, the solution to the case $\beta = 0$ and $m_1 = 0$ is

$$y_{m1}(x) = \begin{cases} m_1 C_{m1,<}\sinh(\alpha x) & \text{if } x < c \\ m_1\left[\cosh(\alpha(1-x)) - 1 + C_{m1,>}\sinh(\alpha(1-x))\right] & \text{if } x > c \end{cases}. \tag{S18}$$

with

$$C_{m1,<} = \frac{1-\cosh(\alpha(1-c))}{\sinh(\alpha)}, \tag{S19}$$

$$C_{m1,>} = \frac{\cosh(\alpha c)}{\sinh(\alpha)} - \coth(\alpha). \tag{S20}$$

As the problem is linear, the full solution of Eqs. (S6) to (S10) is the sum $y(x) = y_\beta(x) + y_{m0}(x) + y_{m1}(x)$. From Eq. (S11), we have $C_2 = \beta C_{\beta,<} + m_0 C_{m_0,<} + m_1 C_{m_1,<}$ and $C_4 = \beta C_{\beta,>} + m_0 C_{m_0,>} + m_1 C_{m_1,>}0$.

In the following, we consider that the beam is clamped at both ends. Equivalently, the derivative of its displacement is null at these positions: $\frac{dy}{dx}(0) = \frac{dy}{dx}(1) = 0$. The two moments $m_0$ and $m_1$ are then determined by this additional constrain. By using Eq. (S11) and with explicit expressions for $C_2$ and $C_4$, we obtain the two following moments

$$m_0 = -\frac{\beta}{\alpha}\frac{(1 - c + \alpha C_{\beta,>})C_{m1,>} - (c + \alpha C_{\beta,>})C_{m1,<}}{C_{m0,<}C_{m1,>} - C_{m0,>}C_{m1,<}}, \tag{S21}$$

$$m_1 = -\frac{\beta}{\alpha}\frac{(1 - c + \alpha C_{\beta,>})C_{m0,>} - (c + \alpha C_{\beta,>})C_{m0,<}}{C_{m1,<}C_{m0,>} - C_{m1,>}C_{m0,<}}. \tag{S22}$$

When the load is applied at the center of the beam $c = 1/2$, these moments are equal by symmetry, and they can be expressed as $m_0 = m_1 = \frac{\beta \tanh(\alpha/4)}{2\alpha}$. The maximum is also located at the center of the beam and is

$$y_{\mathsf{max}} = \frac{\beta}{4}\left[1 - \frac{\tanh(\alpha/4)}{\alpha/4}\right]. \tag{S23}$$

In the limit where $\alpha/4 \gg 1$, we have $y_{\mathsf{max}} = \frac{\beta}{4}$ and $m_0 = m_1 = 0$, which corresponds to $Y_{\mathsf{max}} = \frac{PL}{4T}$. In the limit where $\alpha/4 \ll 1$, we have $y_{\mathsf{max}} = \frac{\beta\alpha^2}{192}$ with the non-zero moments $m_0 = m_1 = \frac{1}{2}\frac{\beta\alpha^2}{192}$, which corresponds to $Y_{\mathsf{max}} = \frac{PL^3}{192EI}$. The apparent stiffness is usually defined as $k_{\mathsf{app}} = \frac{P}{Y_{\mathsf{max}}}$. Using Eq.(S23) with dimensions yields

$$k_{\mathsf{app}} = \frac{4T/L}{1 - \frac{\tanh(L/4\lambda)}{L/4\lambda}}. \tag{S24}$$

In the two limits $\alpha/4 \ll 1$ and $\alpha/4 \gg 1$, we then respectively have $k_{\mathsf{app}} = \frac{4T}{L}$ and $k_{\mathsf{app}} = \frac{192EI}{L^3}$.

#### II. COMPUTATION OF THE STIFFNESS MATRIX

In this section, we compute the stiffness matrix associated with a beam under extensive stress using the formulation of Ref. [2]. The stiffness matrix $[K]$ relates the displacement and tangential angles associated to the forces and moments applied at any position along a beam. We use a finite element model where the beam is discretized into $N$ nodes. Let us focus on the relation between two neighboring nodes that we call 1 and 2 separated by a distance $l = L/(N-1)$ and respectively located at $X = 0$ and $X = l$. For each node we denote $U_i$ the displacement in the positive $Y$-direction, and $\theta_i$ the rotation angle of the beam along the $Z$-axis. We want to find the displacements $U_1$, $U_2$ and rotations $\theta_1$, $\theta_2$ caused by a given set of forces $F_1$, $F_2$ and moments $M_1$, $M_2$ applied at nodes 1, 2. Because we assume that the deflections are small, the rotation angle is given by the local derivative $\theta(x) = \frac{du}{dx}$. In the framework of Finite Element Model (FEM), the distance between nodes is assumed sufficiently small to have a linear relation between the forces and the moments and the displacements and the rotations. Therefore,

$$\{F\} = \begin{pmatrix} F_1 \\ M_1 \\ F_2 \\ M_2 \end{pmatrix} = \begin{pmatrix} K_{11} & K_{12} & K_{13} & K_{14} \\ K_{21} & K_{22} & K_{23} & K_{24} \\ K_{31} & K_{32} & K_{33} & K_{34} \\ K_{41} & K_{42} & K_{43} & K_{44} \end{pmatrix} \begin{pmatrix} U_1 \\ \theta_1 \\ U_2 \\ \theta_2 \end{pmatrix} = [K]\{U\}, \tag{S25}$$

where we have introduced the stiffness matrix $[K]$. Note that the elements of this matrix do not have the same unit. The extension in the longitudinal direction of the beam and the rotation about the beam axis are not considered here.

The continuous beam deflection from node 1 to node 2 is denoted $U(X)$ and we must obey $U(0) = U_1$, $\frac{dU}{dX}(0) = \theta_1$, $U(l) = U_2$, and $\frac{dU}{dX}(l) = \theta_2$. The simplest function for $U(X)$ that is consistent with these boundary conditions is the cubic polynomial $U(X) = a_0 + a_1X + a_2X^2 + a_3X^3$. the coefficients $a_i$ can be computed from the boundary conditions yielding

$$a_0 = U_1, \tag{S26}$$

$$a_1 = \theta_1, \tag{S27}$$

$$a_2 = -3\frac{U_1}{l^2} - 2\frac{\theta_1}{l} + 3\frac{U_2}{l^2} - \frac{\theta_2}{l}, \tag{S28}$$

$$a_3 = 2\frac{U_1}{l^3} + \frac{\theta_1}{l^2} - 2\frac{U_2}{l^3} + \frac{\theta_2}{l^2}. \tag{S29}$$

$$\tag{S30}$$

We can then write that the deformation is

$$U(X) = U_1 + \theta_1 X + \left(-3\frac{U_1}{l^2} - 2\frac{\theta_1}{l} + 3\frac{U_2}{l^2} - \frac{\theta_2}{l}\right) X^2 + \left(2\frac{U_1}{l^3} + \frac{\theta_1}{l^2} - 2\frac{U_2}{l^3} + \frac{\theta_2}{l^2}\right) X^3 \tag{S31}$$

$$= \underbrace{\left(1 - 3\frac{X^2}{l^2} + 2\frac{X^3}{l^3}\right)}_{N_1(X)} U_1 + \underbrace{\left(\frac{X}{l} - 2\frac{X^2}{l^2} + \frac{X^3}{l^3}\right)}_{N_2(X)} l\,\theta_1 + \underbrace{\left(3\frac{X^2}{l^2} - 2\frac{X^3}{l^3}\right)}_{N_3(X)} U_2 + \underbrace{\left(-\frac{X^2}{l^2} + \frac{X^3}{l^3}\right)}_{N_4(X)} l\,\theta_2 \tag{S32}$$

$$= \begin{bmatrix} N_1 & N_2 & N_3 & N_4 \end{bmatrix} \begin{bmatrix} U_1 \\ \theta_1 \\ U_2 \\ \theta_2 \end{bmatrix} \tag{S33}$$

$$= [N]\{U\}. \tag{S34}$$

Let us note that the $X$-dependency of $U$ is now contained in $[N]$.

The Castigliano's theorem states that the force $F_i$ (resp. the moment $M_i$) that is exerted on node $i$ is the derivative of the total energy $\mathcal{E}_{\text{tot}}$ with respect to the displacement $U_i$ (resp. the rotation $\theta_i$) at node $i$. Here, we consider two contributions to the total energy. One is the strain energy $\mathcal{E}_b$ that is related to the bending rigidity of the beam, while the other is the energy $\mathcal{E}_s$ that is related to the extensive stress applied at both ends of the beam. From Castigliano's theorem, we have

$$\{F\} = \nabla\mathcal{E}_{\text{tot}} = \nabla(\mathcal{E}_b + \mathcal{E}_s) = ([K_b] + [K_s])\{U\} = [K]\{U\}, \tag{S35}$$

where we have introduced the operator $\nabla = (\frac{\partial}{\partial U_1}\frac{\partial}{\partial \theta_1}\frac{\partial}{\partial U_2}\frac{\partial}{\partial \theta_2})$ as well as the stiffness matrices $[K_b]$ and $[K_s]$, which will be defined later. Let us focus on the strain energy $\mathcal{E}_b$ first. The strain energy can be explicitly computed as (see Chap. 4 of Ref. [2])

$$\mathcal{E}_b = \frac{1}{2}\int dV E Y^2 \left(\frac{d^2U}{dX^2}\right)^2, \tag{S36}$$

$$= \frac{1}{2}E\underbrace{\left(\int Y^2 dA\right)}_{=I}\int_0^l \left(\frac{d^2[N]}{dX^2}\{U\}\right)^2 dX, \tag{S37}$$

$$= \frac{1}{2}EI\int_0^l \left(\frac{d^2N_1}{dX^2}U_1 + \frac{d^2N_2}{dX^2}\theta_1 + \frac{d^2N_3}{dX^2}U_2 + \frac{d^2N_4}{dX^2}\theta_2\right)^2 dX, \tag{S38}$$

where $E$ is the Young modulus of the beam, and $I$ is its moment of inertia. This gives

$$F_1 = \frac{\partial \mathcal{E}_b}{\partial U_1} = EI\int_0^l \left(\frac{d^2N_1}{dX^2}U_1 + \frac{d^2N_2}{dX^2}\theta_1 + \frac{d^2N_3}{dX^2}U_2 + \frac{d^2N_4}{dX^2}\theta_2\right)\frac{d^2N_1}{dX^2}dX. \tag{S39}$$

This expression can be related to Eq. (S25) and provides a way to compute the elements of the stiffness matrix $K_{1i}$. As an example, let us compute the element $K_{11}$ appearing in Eq. (S25)

$$K_{11} = EI\int_0^l \left(\frac{d^2N_1}{dX^2}\right)^2 dX \tag{S40}$$

$$= \frac{EI}{l^3}\int_0^l \left(-6 + 12\frac{X}{l}\right)^2 \frac{dX}{l} \tag{S41}$$

$$= 12EI. \tag{S42}$$

Computing systematically the other elements $K_{ii}$ of the bending stiffness matrix yields

$$[K_b] = \frac{EI}{l^3}\begin{pmatrix} 12 & 6l & -12 & 6l \\ 6l & 4l^2 & -6l & 2l^2 \\ -12 & -6l & 12 & -6l \\ 6l & 2l^2 & -6l & 4l^2 \end{pmatrix} \tag{S43}$$

Let us now turn to the potential energy $\mathcal{E}_s$ related to the extensive stress $T$ applied at both ends of the beam. Using the expression found in Ref. [3], we have

$$\mathcal{E}_s = \frac{1}{2}\int_0^l T\left(\frac{dU}{dX}\right)^2 dX \tag{S44}$$

$$= \frac{1}{2}T\int_0^l \left(\frac{dN_1}{dX}U_1 + \frac{dN_2}{dX}\theta_1 + \frac{dN_3}{dX}U_2 + \frac{dN_4}{dX}\theta_2\right)^2 dX. \tag{S45}$$

To illustrate how to compute the stiffness matrix related to the extensive stress, we compute the moment $M_1$ applied on node 1. Using Castigliano's theorem

$$M_1 = T\int_0^l \left(\frac{dN_1}{dX}U_1 + \frac{dN_2}{dX}\theta_1 + \frac{dN_3}{dX}U_2 + \frac{dN_4}{dX}\theta_2\right)\frac{dN_2}{dX}dX. \tag{S46}$$

This expression can be related to Eq. (S25) and provides a way to compute the elements of the stiffness matrix $K_{2i}$. As an example, let us compute the element $K_{21}$ appearing in Eq. (S25)

$$K_{21} = T\int_0^l \frac{dN_1}{dX}\frac{dN_2}{dX}dX \tag{S47}$$

$$= T\int_0^l \left(-6\frac{X}{l}+6\frac{X^2}{l^2}\right)\left(1-4\frac{X}{l}+3\frac{X^2}{l^2}\right)dx \tag{S48}$$

$$= \frac{T}{10} \tag{S49}$$

Computing the other elements $K_{ii}$ of the bending stiffness matrix yields

$$[K_s] = \frac{T}{10l}\begin{pmatrix} 12 & l & -12 & l \\ l & \frac{4l^2}{3} & -l & -\frac{l^2}{3} \\ -12 & -l & 12 & -l \\ l & -\frac{l^2}{3} & -l & \frac{4l^2}{3} \end{pmatrix}. \tag{S50}$$

With the expressions of the stiffness matrices, we can rewrite Eq. (S35) in a dimensionalized form as

$$\{F\} = \begin{pmatrix} F_1 \\ M_1 \\ F_2 \\ M_2 \end{pmatrix} = \left(\frac{EI}{l^3}\begin{pmatrix} 12 & 6l & -12 & 6l \\ 6l & 4l^2 & -6l & 2l^2 \\ -12 & -6l & 12 & -6l \\ 6l & 2l^2 & -6l & 4l^2 \end{pmatrix} + \frac{T}{10l}\begin{pmatrix} 12 & l & -12 & l \\ l & \frac{4l^2}{3} & -l & -\frac{l^2}{3} \\ -12 & -l & 12 & -l \\ l & -\frac{l^2}{3} & -l & \frac{4l^2}{3} \end{pmatrix}\right)\begin{pmatrix} U_1 \\ \theta_1 \\ U_2 \\ \theta_2 \end{pmatrix} \tag{S51}$$

Let us extend this analysis to both $Y$ and $Z$ displacements of the beam. We introduce the displacement $U_i^z$ in the $Z$-direction and its associated rotational angle $\theta_i^y$ about the $Y$-axis for each node $i$. Conversely, the displacement in the $Y$-direction is called $U_i^y$ and its associated rotational angle about the $Z$-axis $\theta_i^z$. The name of the forces and moments applied at each node $i$ are changed accordingly. We define the stiffness matrices then obey

$$\{F\} = \begin{pmatrix} F_1^y \\ M_1^z \\ F_2^y \\ M_2^z \\ F_1^z \\ M_1^y \\ F_2^z \\ M_2^y \end{pmatrix} = \left([K_b^{2d}] + [K_s^{2d}]\right)\begin{pmatrix} U_1^y \\ \theta_1^z \\ U_2^y \\ \theta_2^z \\ U_1^z \\ \theta_1^y \\ U_2^z \\ \theta_2^y \end{pmatrix} = ([K_b^{2d}] + [K_s^{2d}])\{U\} \tag{S52}$$

We assume that motions in $(U_i^y, \theta_i^z)$ are decoupled from motions in $(U_i^z, \theta_i^y)$. Therefore,

$$[K_i^{2d}] = \begin{pmatrix} [K_i^{xy}] & [0] \\ [0] & [K_i^{xz}] \end{pmatrix}, \tag{S53}$$

where $i = b, s$, and $[0]$ is a $(4 \times 4)$ matrix with all components set to zero. Above, we computed the stiffness matrices for motions in the $(X, Y)$-plane, which holds true here: $[K_b^{xy}] = [K_b]$ and $[K_s^{xy}] = [K_s]$, with $[K_b]$ and $[K_s]$ respectively defined in Eqs. (S43) and (S50). To compute the stiffness matrices $[K_b^{xz}]$ and $[K_s^{xz}]$ for motions in the $(X, Z)$-plane, we must adapt the above reasoning to our sign convention for the rotation angles. All the angles are defined positive for anticlockwise rotation about their reference axis, e.g. $\theta_1^z$ is positive for anti-clockwise rotation of the beam about the $Z$-axis at node 1. With right-handed convention for the $(X, Y, Z)$ reference frame, if the longitudinal direction of the beam is along the positive $X$-axis, we have $\theta^z = \frac{dU^y}{dX}$, and $\theta^y = -\frac{dU^z}{dX}$. Therefore, the reasoning detailed above for $(U_i^y, \theta_i^z)$ must be changed for the motions in $(U_i^z, \theta_i^y)$ by using the transformation $(U_i^z, \theta_i^y) \to (U_i^z, -\theta_i^y)$ and $(F_i^z, M_i^y) \to (F_i^z, -M_i^y)$. With this transformation, we find that the stiffness matrix related to the bending rigidity is

$$[K_b^{2d}] = \frac{EI}{l^3}\begin{pmatrix} 12 & 6l & -12 & 6l & 0 & 0 & 0 & 0 \\ 6l & 4l^2 & -6l & 2l^2 & 0 & 0 & 0 & 0 \\ -12 & -6l & 12 & -6l & 0 & 0 & 0 & 0 \\ 6l & 2l^2 & -6l & 4l^2 & 0 & 0 & 0 & 0 \\ 0 & 0 & 0 & 0 & 12 & -6l & -12 & -6l \\ 0 & 0 & 0 & 0 & -6l & 4l^2 & 6l & 2l^2 \\ 0 & 0 & 0 & 0 & -12 & 6l & 12 & 6l \\ 0 & 0 & 0 & 0 & -6l & 2l^2 & 6l & 4l^2 \end{pmatrix}. \tag{S54}$$

By convention, the vectors $\{F\}$ and $\{U\}$ are usually shown in the following order $\{F\} = (F_1^y, F_1^z, M_1^y, M_1^z, F_2^y, F_2^z, M_2^y, M_2^z)^T$ and $\{U\} = (U_1^y, U_1^z, \theta_1^y, \theta_1^z, U_2^y, U_2^z, \theta_2^y, \theta_2^z)^T$. With , $[K_b^{2d}]$ becomes

$$[K_b^{2d}] = \frac{EI}{l^3}\begin{pmatrix} 12 & 0 & 0 & 6l & -12 & 0 & 0 & 6l \\ 0 & 12 & -6l & 0 & 0 & -12 & -6l & 0 \\ 0 & -6l & 4l^2 & 0 & 0 & 6l & 2l^2 & 0 \\ 6l & 0 & 0 & 4l^2 & -6l & 0 & 0 & 2l^2 \\ -12 & 0 & 0 & -6l & 12 & 0 & 0 & -6l \\ 0 & -12 & 6l & 0 & 0 & 12 & 6l & 0 \\ 0 & -6l & 2l^2 & 0 & 0 & 6l & 4l^2 & 0 \\ 6l & 0 & 0 & 2l^2 & -6l & 0 & 0 & 4l^2 \end{pmatrix}. \tag{S55}$$

Applying the same transformation to the stiffness matrix $[K_s^{2d}]$ that is related to the stress applied at both ends of the beam yields

$$[K_s^{2d}] = \frac{P}{10l}\begin{pmatrix} 12 & 0 & 0 & l & -12 & 0 & 0 & l \\ 0 & 12 & -l & 0 & 0 & -12 & -l & 0 \\ 0 & -l & \frac{4l^2}{3} & 0 & 0 & l & -\frac{l^2}{3} & 0 \\ l & 0 & 0 & \frac{4l^2}{3} & -l & 0 & 0 & -\frac{l^2}{3} \\ -12 & 0 & 0 & -l & 12 & 0 & 0 & -l \\ 0 & -12 & l & 0 & 0 & 12 & l & 0 \\ 0 & -l & -\frac{l^2}{3} & 0 & 0 & l & \frac{4l^2}{3} & 0 \\ l & 0 & 0 & -\frac{l^2}{3} & -l & 0 & 0 & \frac{4l^2}{3} \end{pmatrix}. \tag{S56}$$

We have calculated the matrices that describe the displacement of two neighboring nodes along the beam in response to forces and moments applied at each of those two nodes. These matrices can be extended to any pair of neighboring nodes along the beam. This provides a matrix that is diagonal by

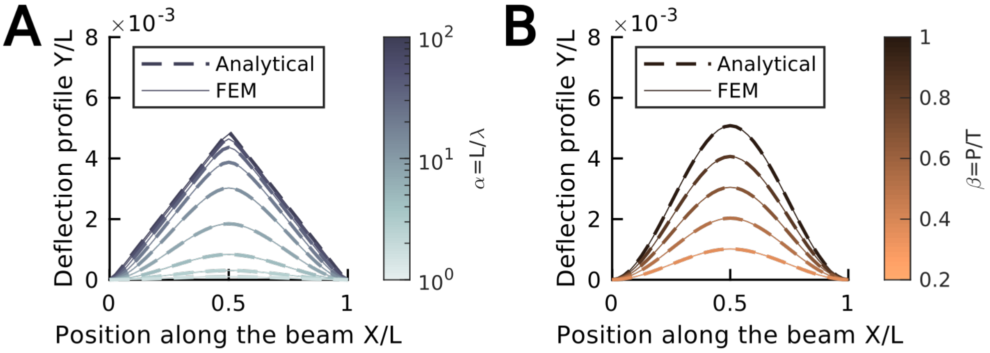

Sup.Fig.N1.2: Comparison of the displacement of a clamped beam in response to a point load $P$ applied at $C = L/2$ obtained with the analytical (Section I, dashed line) and FEM (Section II, continuous line) calculation. The deflection profiles $Y/L$ are plotted as a function of the normalized position $X/L$ along the beam for varying adimensional parameters: **A**: $\beta = 0.02$, and $\alpha$ is varied logarithmically from $1$ to $100$ with a multiplicative factor of $10^{1/4}$, **B**: $\beta$ is varied linearly from $0.2$ to $1$ with an increment of $0.2$ and $\alpha = 1$. The beam was discretized into $41$ nodes for all FEM calculations.

block, and that can describe the full beam if appropriate boundary conditions are provided at the beam ends.

#### III. VALIDATION OF THE FINITE ELEMENT MODEL

To validate the Finite Element Model (FEM) developed in Section II, we confront it to the analytical solution found in Section I for the case of a point load $P$ applied at the center of the beam ($C = L/2$). Both ends of the beam are assumed to be clamped in the FEM model, matching the boundary conditions presented in Section I. The problem then depends only on the two adimensional parameters $\alpha$ and $\beta$ defined in Section I (the third one $c$ corresponds to the point where the load is applied and is here fixed to $c = 1/2$). To ensure a fast computational time ($< 10$ ms per run) and a small relative error ($< 3\%$) of the FEM calculation with respect to the analytical calculation, we choose to discretize the beam into $41$ nodes. To describe the full beam using FEM, we adapted the package calfem (https://github.com/CALFEM/calfem-matlab) by changing the stiffness matrix of the beam3e function to the sum of the stiffness matrices in Eqs. (S55) and (S56). Sup.Fig.N1.2 shows that the FEM calculation matches the analytical one for any value of $\alpha$ (representative values shown on Sup.Fig.N1.2A, and any value of $\beta$ (representative values shown on Sup.Fig.N1.2B).

---

[1] S. Timoshenko, *Strength of Materials. 2: Advanced Theory and Problems* (Van Nostrand, 1956).
[2] D. Hutton, *Fundamentals of finite element analysis* (McGraw-Hill, 2004).
[3] B. Tu-Sekine, A. Padhi, S. Jin, S. Kalyan, K. Singh, M. Apperson, R. Kapania, S. C. Hur, A. Nain, and S. F. Kim, The FASEB Journal **33**, 14137 (2019).

### Supplementary Note 2 - Evaluation of the force reconstruction optimization workflow and choice of regularization parameters

#### I. Unconstrained two-area loading without noise

A common situation observed in our setup involved an elongated cell fully spread across several fibers, exerting two poles of traction forces at each end of the contacted fibers. Traction force recovery in this situation is more challenging than in the case of a localized loading exerted by a local protrusion, where we constrain the traction localization to a limited number of nodes at the intersection between the cell body and the fiber. Therefore, we chose this limiting case to optimize regularization parameters and assess the performance of our force reconstruction process.

In our simulation, an 80 µm-long fiber was divided into 41 nodes, and a 20 nN load was uniformly distributed along the *y* and *z* axes across two regions of 3 nodes each at *L*/6 and 5*L*/6 (Sup.Fig.N2.1A). We generated the corresponding deflection profiles and we used them to recover the applied forces using our optimization scheme.

##### 1. Choice of *Ɛ*

The first parameter to set is *Ɛ*, the step size of the cost function gradient used in the optimization process:

$$\frac{\partial \mathrm{C}(\mathrm{p_0}, \dots, \mathrm{p_i}, \dots, \mathrm{p_{Nb\_nodes}})}{\partial \mathrm{p_i}} = \frac{\mathrm{C}\big(\mathrm{p_0}, \dots, \mathrm{p_i} + \mathrm{Ɛ}, \dots, \mathrm{p_{Nb\_nodes}}\big) - \mathrm{C}\big(\mathrm{p_0}, \dots, \mathrm{p_i}, \dots, \mathrm{p_{Nb\_nodes}}\big)}{\mathrm{Ɛ}}$$

where $C$ is the optimization cost function and $p$ is the load vector.
Larger values of *Ɛ* fail to converge to the expected solution while smaller values lead to larger computation time. Sup.Fig.N2.2A, B illustrate the force profiles obtained for different values of *Ɛ*.

We define two error metrics (Sup.Fig. N2.1B, C):

- **Contractility error**, accounts for the deviation in total absolute value of the exerted forces between the simulation and the reconstruction:

$$\boldsymbol{Err_c} = \frac{\left|\sum_{k=1}^{Nb\_nodes} \left(|p^{true}{}_{k,y}| + |p^{true}{}_{k,z}|\right) - \sum_{k=1}^{Nb\_nodes} \left(|p^{reconstructed}{}_{k,y}| + |p^{reconstructed}{}_{k,z}|\right)\right|}{\sum_{k=1}^{Nb\_nodes} \left(|p^{true}{}_{k,y}| + |p^{true}{}_{k,z}|\right)}$$

- **Force profile error,** defined as the sum of the absolute deviation in force at each node, which penalizes both incorrect force intensity and incorrect force localization:

$$\boldsymbol{Err_{fp}} = \frac{\sum_{k=1}^{Nb\_nodes} \left(|p^{true}{}_{k,y} - p^{reconstructed}{}_{k,y}| + |p^{true}{}_{k,z} - p^{reconstructed}{}_{k,z}|\right)}{\sum_{k=1}^{Nb\_nodes} \left(|p^{true}{}_{k,y}| + |p^{true}{}_{k,z}|\right)}$$

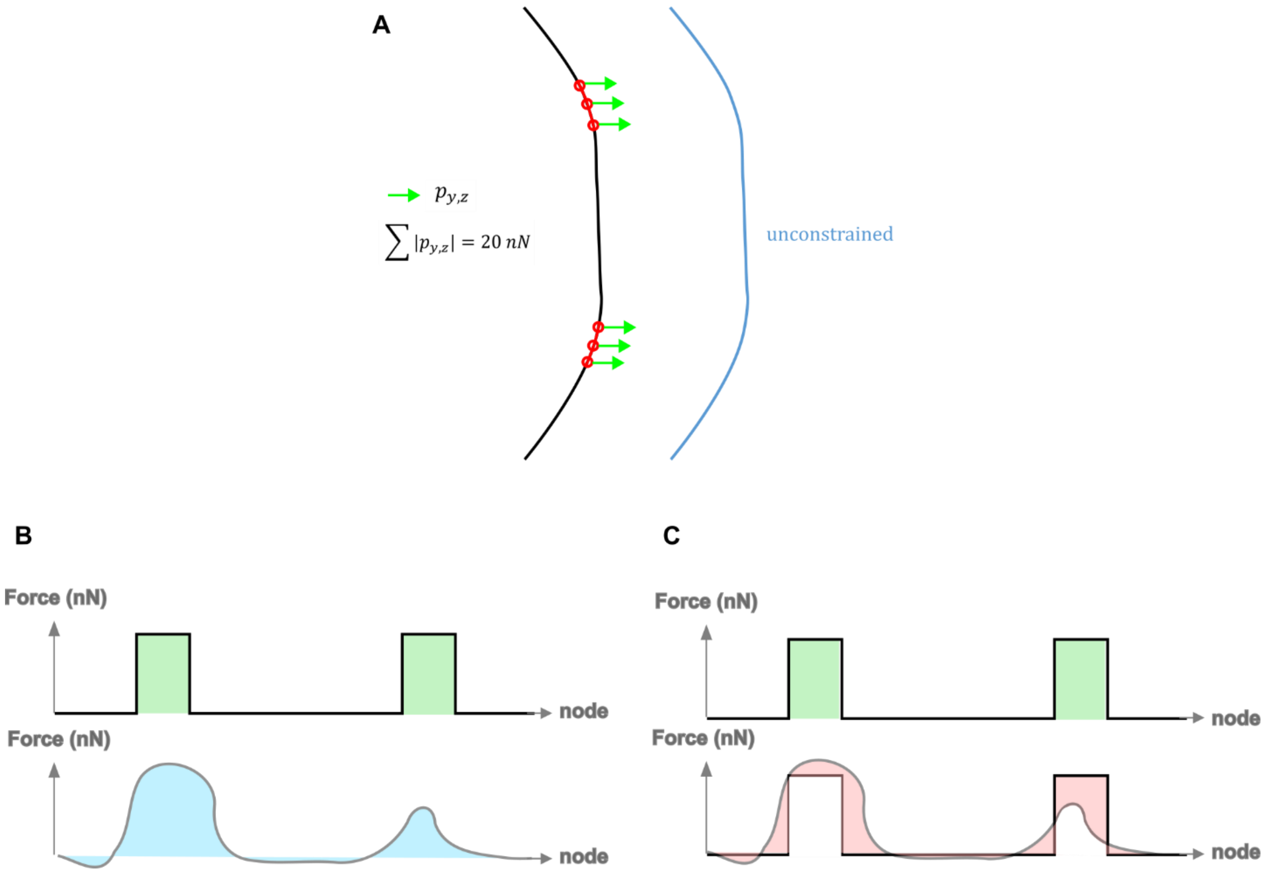

**Supplementary Figure N2.1 - Simulation and error metrics**

**A.** Simulated unconstrained two-area loading. **B.** The contractility error metric compares the simulated contractility (top, green area) and the reconstructed one (bottom, blue). **C.** The force profile error metric evaluates the local underestimation or overestimation of the reconstructed force (bottom, red area).

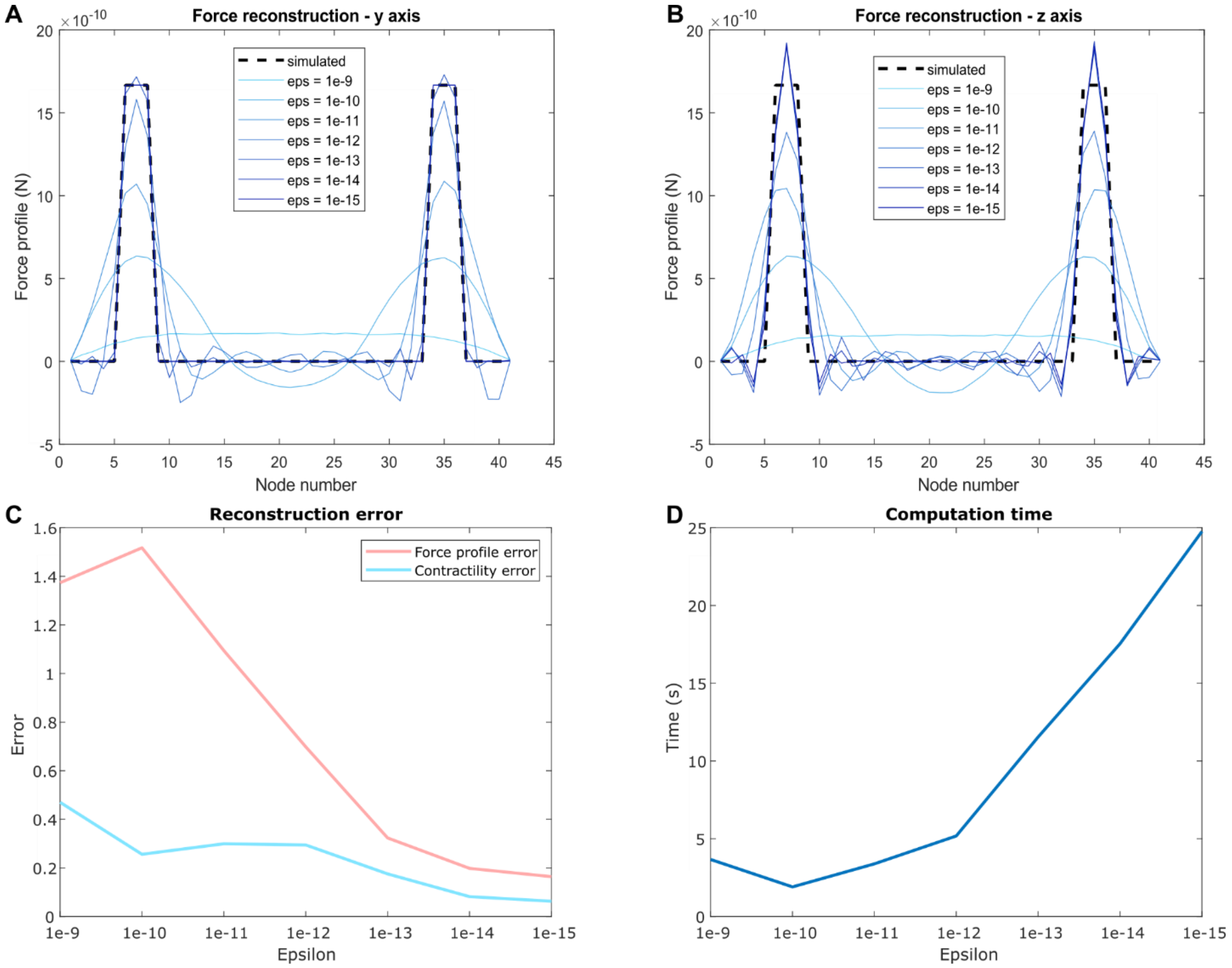

**Supplementary Figure N2.2 – Optimization of the gradient step size** $\mathcal{E}$

**A.** Simulated force profile and corresponding y component and **B.** z component of the reconstructions for different values of the gradient step size $\mathcal{E}$. **C.** Reconstruction errors and **D.** computation time for the force recovery process for different $\mathcal{E}$ values.

Based on the observation of these reconstruction errors and of the computation time, we chose to set $\mathcal{E} = 10^{-13}$ N.

#### 2. Influence of the number of nodes on the reconstruction

In this section, we aim to evaluate how the number of nodes chosen in the Finite Element discretization of fibers affects the performance of the traction force recovery. We simulated a loading pattern similar to the one described in section 1, where the loading area had a fixed length of 6 µm. We generated the corresponding "ideal" deflection profile using a large number of nodes (Nb_nodes = 201). This deflection profile was then subsampled to a lower number of nodes, and force reconstruction was performed using $\mathcal{E} = 10^{-13}$ N. The number of nodes was varied between 5 and 77 (Sup.Fig.N2.3A, B).

For each case, we computed the contractility error and force profile error (Sup.Fig.N2.3C) as well as the computation time (Sup.Fig.N2.3D). We confirmed our choice of Nb_nodes = 41 as a good compromise between reconstruction accuracy and computation time, achieving an excellent agreement between the simulated and reconstructed deflection profiles (Sup.Fig.N2.3E, F). It is worth noting that due to the cross-section anisotropy of the fiber, the moment of inertia differ along the *y* and *z* axes. Although the contribution of bending stiffness to the overall deflection is minor compared to tension, this leads to slightly different deflection profiles along the *y* and *z* directions.

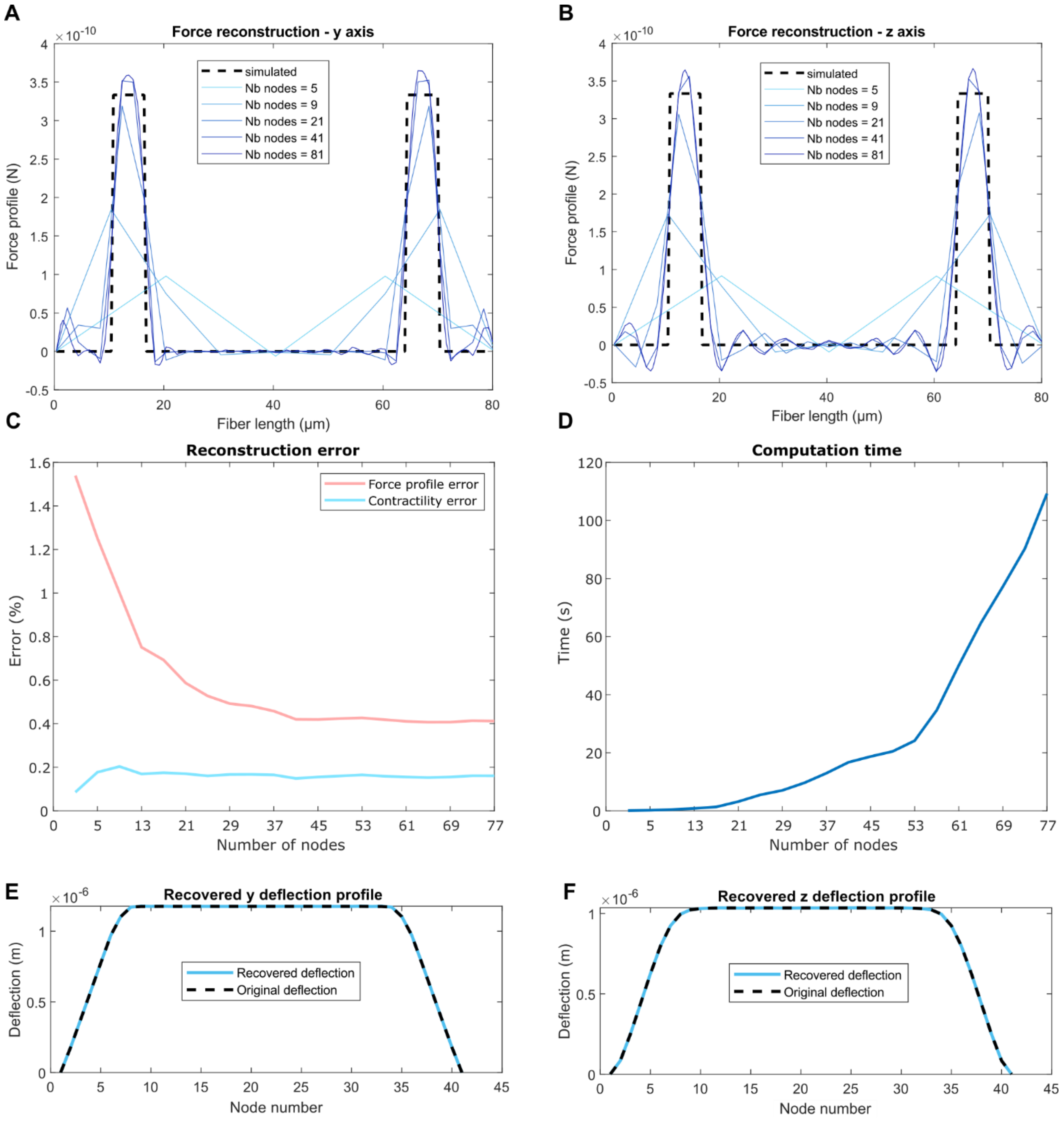

**Supplementary Figure N2.3 – Influence of the number of nodes on the reconstruction**

**A.** Simulated force profile and corresponding y component and **B.** z component of the reconstructions for different number of nodes. **C.** Reconstruction errors and **D.** computation time for the force recovery process for different number of nodes. **E.** Deflection profile corresponding to a reconstruction with $\mathcal{E} = 10^{-13}$ N and 41 nodes along the y direction and **F.** along the z direction.

#### II. Unconstrained two-area loading with noise - Regularization parameters

Unlike the ideal simulated case previously described, the deflection measurements extracted from our system exhibit some noise. The typical standard deviation is 0.3 µm, which corresponds to our final voxel size. To simulate the effect of this noise, we added a Gaussian noise ($\sigma$ = 0.3 µm) to the simulated deflection profile before applying the inverse method for traction recovery. In this scenario, the recovered deflection profile clearly shows overfitting to the added noise (Sup.Fig.N2.4C, D) resulting in poor force reconstruction (Sup.Fig.N2.4A,B).

To adress this, we applied an elastic net regularization (L1+L2) to prevent overfitting. To select optimal regularization parameters $\lambda_1$ and $\lambda_2$, we ran the simulation multiple times (N=10) with various parameter pairs and calculated the average reconstruction errors (Sup.Fig.N2.4E, F). We identified the optimal pair ($\lambda_1=10^2$ m.N$^{-1}$, $\lambda_2=10^9$ m.N$^{-2}$) which minimized both errors, and used these parameters for all regularized reconstructions.

Examples of reconstructed force profiles with regularization are shown in Sup.Fig. N2.4G, H, along with the corresponding deflection profiles (Sup.Fig. N2.4I, J). While the reconstruction is significantly improved, these examples examples also illustrate some limitations encountered in this challenging case, including underestimation or overestimation of local force intensity and shifting of the force peak localization.

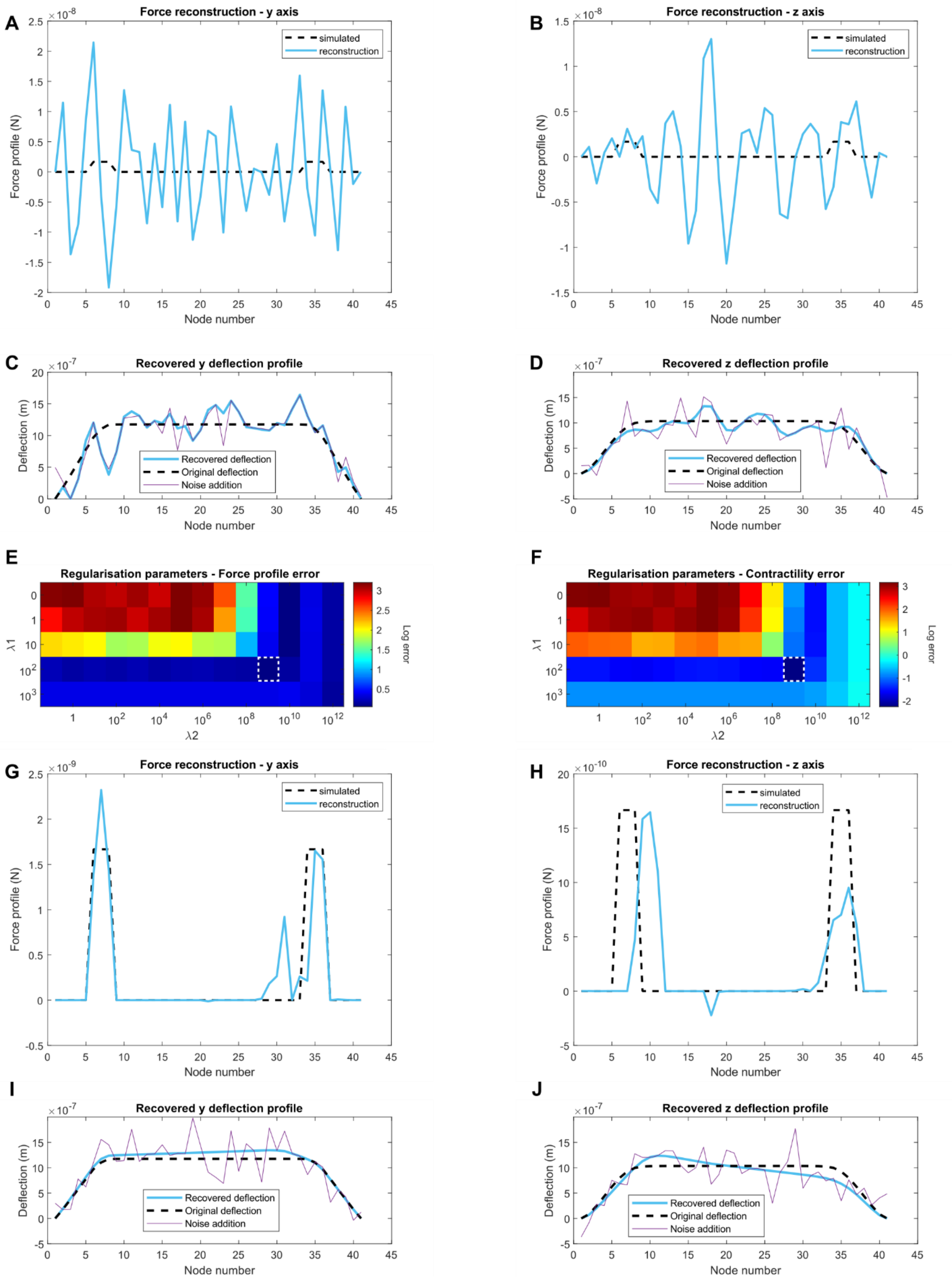
A
Force reconstruction - y axis
simulated
reconstruction
Force profile (N)
Node number
B
Force reconstruction - z axis
simulated
reconstruction
Force profile (N)
Node number
C
Recovered y deflection profile
Recovered deflection
Original deflection
Noise addition
Deflection (m)
Node number
D
Recovered z deflection profile
Recovered deflection
Original deflection
Noise addition
Deflection (m)
Node number
E
Regularisation parameters - Force profile error
λ1
λ2
Log error
F
Regularisation parameters - Contractility error
λ1
λ2
Log error
G
Force reconstruction - y axis
simulated
reconstruction
Force profile (N)
Node number
H
Force reconstruction - z axis
simulated
reconstruction
Force profile (N)
Node number
I
Recovered y deflection profile
Recovered deflection
Original deflection
Noise addition
Deflection (m)
Node number
J
Recovered z deflection profile
Recovered deflection
Original deflection
Noise addition
Deflection (m)
Node number

**Supplementary Figure N2.4 - Unconstrained two-area loading with noise**

**A.** Simulated force profile and corresponding y component and **B.** z component of the reconstructions with addition of a gaussian noise (σ = 0.3 µm) to the simulated deflection profile before the traction recovery process. **C.** Corresponding reconstructed deflection profiles along the y direction and **D.** along the z direction. **E.** Logarithmic representation of the force profile error and **F.** of the contractility error depending on the regularization parameters $\lambda_1$ and $\lambda_2$. The white dashed line square figures the pair of parameters that minimizes both the two reconstruction errors. **G.** Y component of the reconstructed force profile and **H.** z component after regularization. **I.** Corresponding reconstructed deflection profile along the y direction and **J.** along the z direction.

#### III. Constrained one-area loading with noise

To assess the performance of our force reconstruction framework in the case of a localized loading – such as an individual protrusion pulling on a fiber - we simulated a 20 nN loading distributed along the *y* and *z* axes and applied at a single node, located at *L*/4. A Gaussian noise (*σ* = 0.3 µm) was added to the simulated deflection profile. The regularized reconstruction of both the force and deflection profiles are shown in Sup.Fig. N2.5B, C and D,E, demonstrating excellent agreement between the simulation and the reconstruction in this simpler scenario.

We then evaluated the impact of the loading area on reconstruction performance by distributing the 20 nN load over an increasingly larger area, centered around *L*/4. The corresponding reconstruction errors and computation times are displayed in Sup.Fig.N2.5F, G.

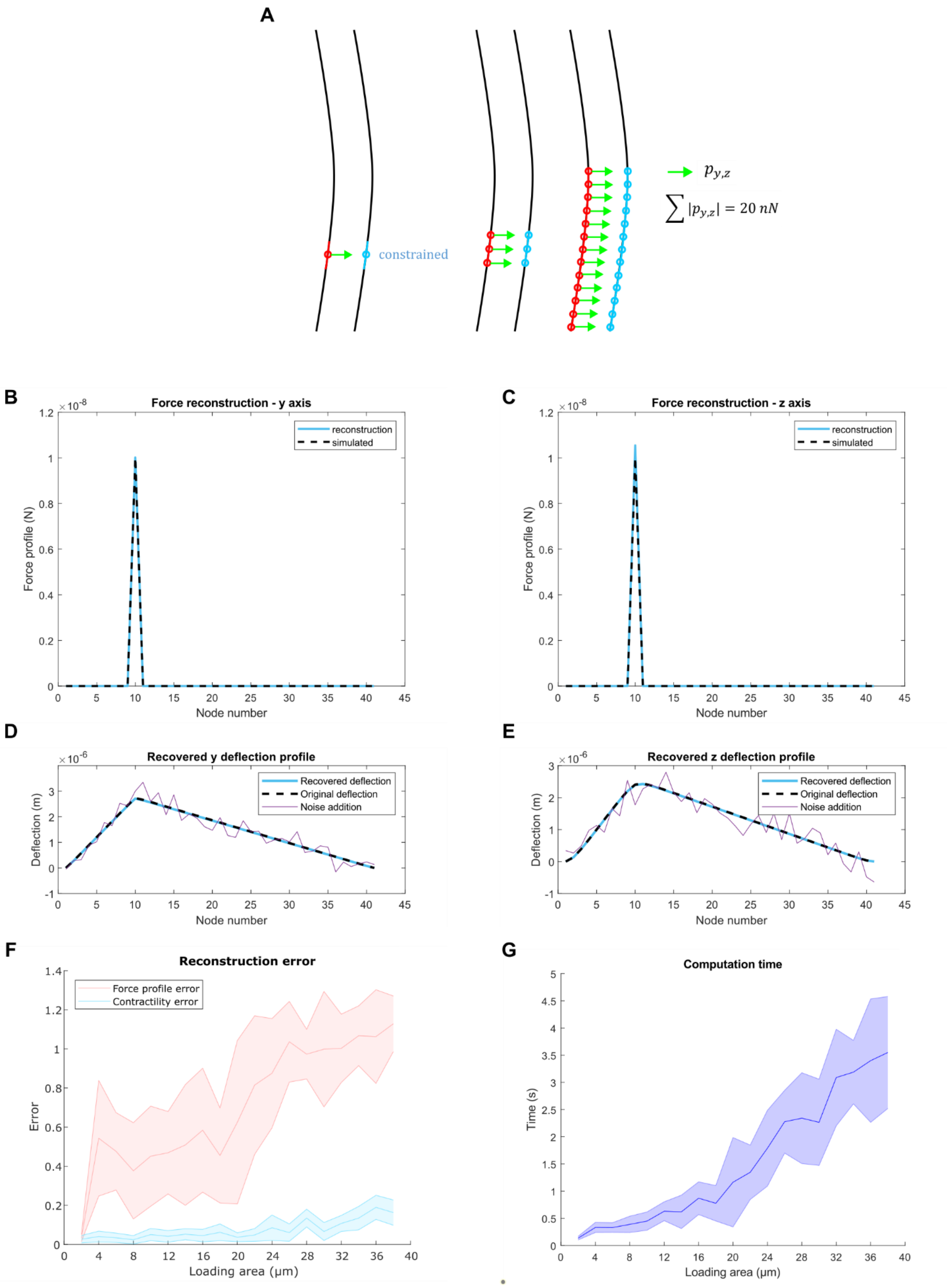
A
constrained
$p_{y,z}$
$\sum |p_{y,z}| = 20\ nN$
B
Force reconstruction - y axis
reconstruction
simulated
Force profile (N)
Node number
C
Force reconstruction - z axis
reconstruction
simulated
Force profile (N)
Node number
D
Recovered y deflection profile
Recovered deflection
Original deflection
Noise addition
Deflection (m)
Node number
E
Recovered z deflection profile
Recovered deflection
Original deflection
Noise addition
Deflection (m)
Node number
F
Reconstruction error
Force profile error
Contractility error
Error
Loading area (μm)
G
Computation time
Time (s)
Loading area (μm)

**Supplementary Figure N2.5 - Constrained one-area loading with noise**

**A.** Simulated constrained one-area loading distributed over an increasing number of nodes. **B.** Simulated force profile and corresponding y component and **C.** z component of the reconstructions for a localized loading applied at L/4. **D.** Corresponding reconstructed deflection profiles along the y direction and **E.** along the z direction. **F.** Reconstruction errors and **G.** computation time of the traction recovery process as a function of the loading area (1 node = 2 µm) for a constant total contractility of 20 nN.

#### Supplementary Note 3 - Residual forces allow to evaluate the underestimation of traction forces due to neglecting the axial component

Cell traction forces should sum to zero. We quantified the residual forces, defined as the absolute value of the summed x-y components $|\sum p_{x,y}|$ or z component $|\sum p_z|$ of the traction forces at a given time point. We observed that when a cell contacts fibers on the two orthogonal planes, the x-y component of the residual forces often peaks, indicating an apparent force imbalance due to neglecting the axial component. In this case (Sup.Fig.N3.1), the bending of fibers in one plane is counterbalanced by an axial force exerted on the orthogonal layer. Analyzing the residual force components allows us to estimate the contribution of axial forces in certain cases. For example, in the scenario shown in Sup.Fig.N3.1 ($T_1$) , this contribution reached up to 65% of the measured contractility during contractile bursts. During phases where the cell contacted only one fiber layer ($T_2$), the x-y component of the residual forces remained close to zero. It is important to note that this approach does not account for axial forces balanced within each independent fiber layer.

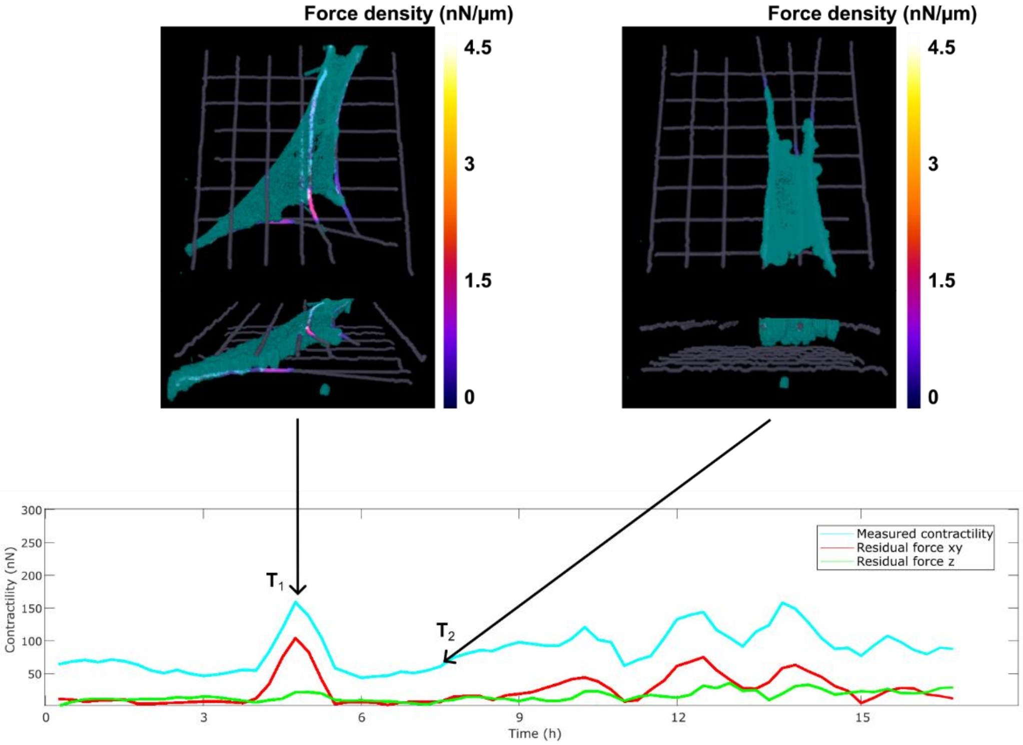

**Supplementary Figure N3.1 - Residual forces along x-y and along z**

Quantification of the x-y and z components of the residual forces for a HUVEC on a 2-layer fiber array with 10 µm lateral spacing and low-exposure fibers ($T_{exp}$ = 600 µs, k = 3.5 nN/µm). The x-y component peaks at $T_1$, when the cell is spread on the two levels of fibers. By contrast, it remains close to zero at $T_2$, when the cell is only spread on the top layer.

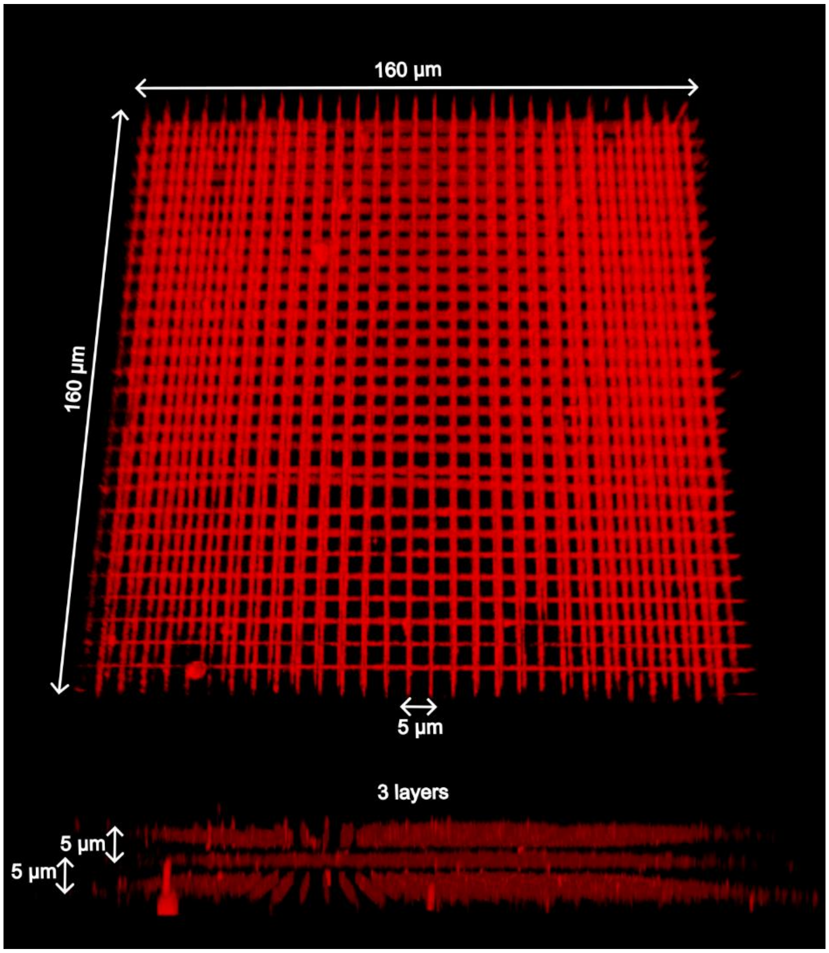

**Supplementary Figure 1 – Multilayer 3D scaffolds**

3D view of a 3-layer 3D microscaffold with both lateral spacing and interlayer spacing set to 5 µm (top: top view, bottom: side view; red: fluorescent fibronectin coated fibers, spinning disk imaging).

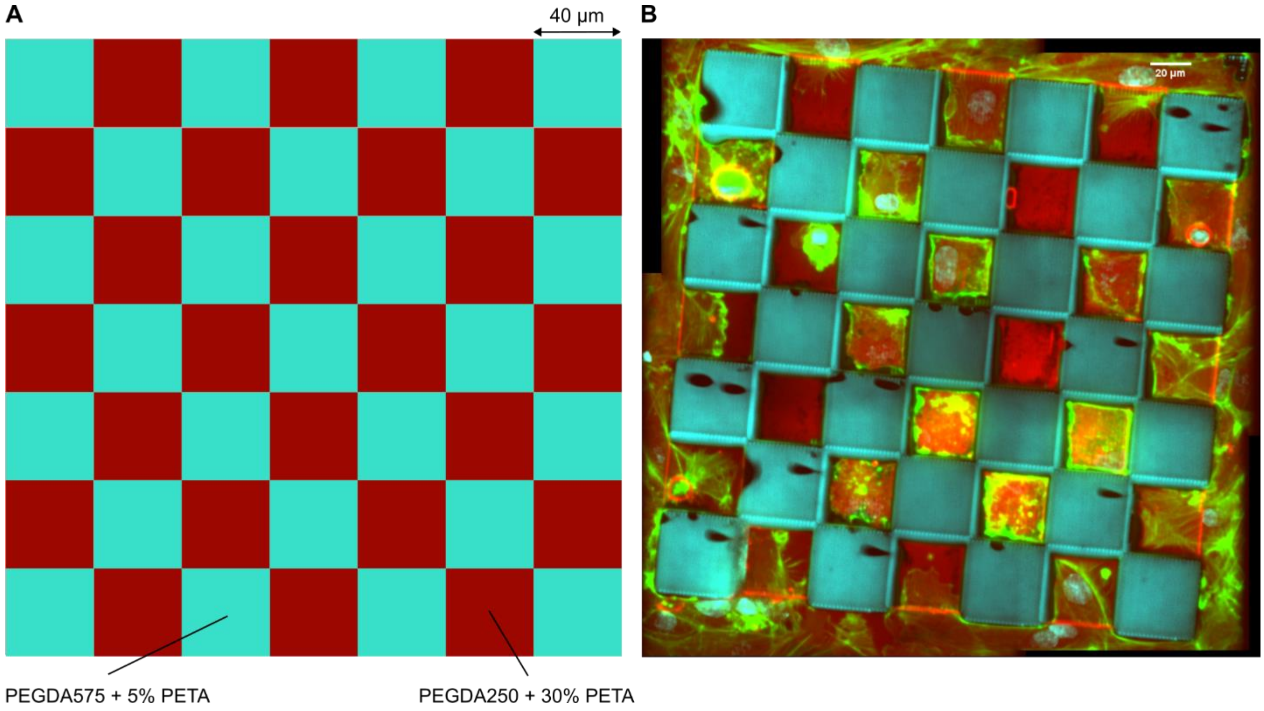

**Supplementary Figure 2 - Spatially directed adhesion in composite microstructures**

**A.** Schematic of a composite checkerboard 2D microstructure used to evaluate the choice of materials used to direct protein adsorption and cell adhesion. **B.** Actin (green) and nuclei (blue) staining of HUVEC cells seeded at high density on a fluorescent fibronectin (red) coated composite checkerboard and fixed after two hours. Cells exclusively adhere to the PEGDA250 part and adopt a square shape.

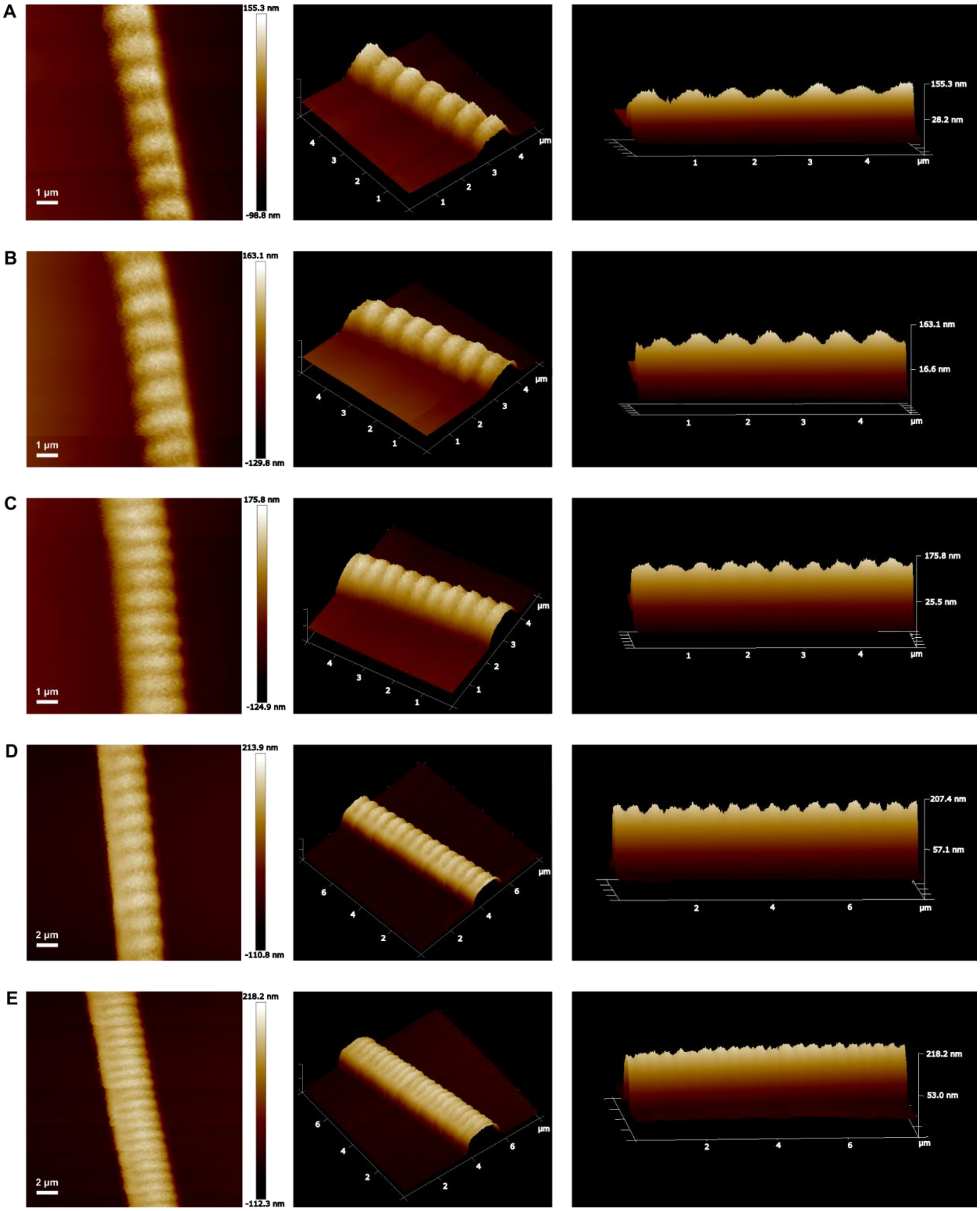

**Supplementary Figure 3 - AFM topography scanning of the photopolymerized fibers**

Topography images (left) and 3D views (middle: top view, right: side view) of the fibers for **A.** $T_{exp}$ = 600 µs, **B.** $T_{exp}$ = 650 µs, **C.** $T_{exp}$ = 700 µs, **D.** $T_{exp}$ = 750 µs and **E.** $T_{exp}$ = 800 µs.

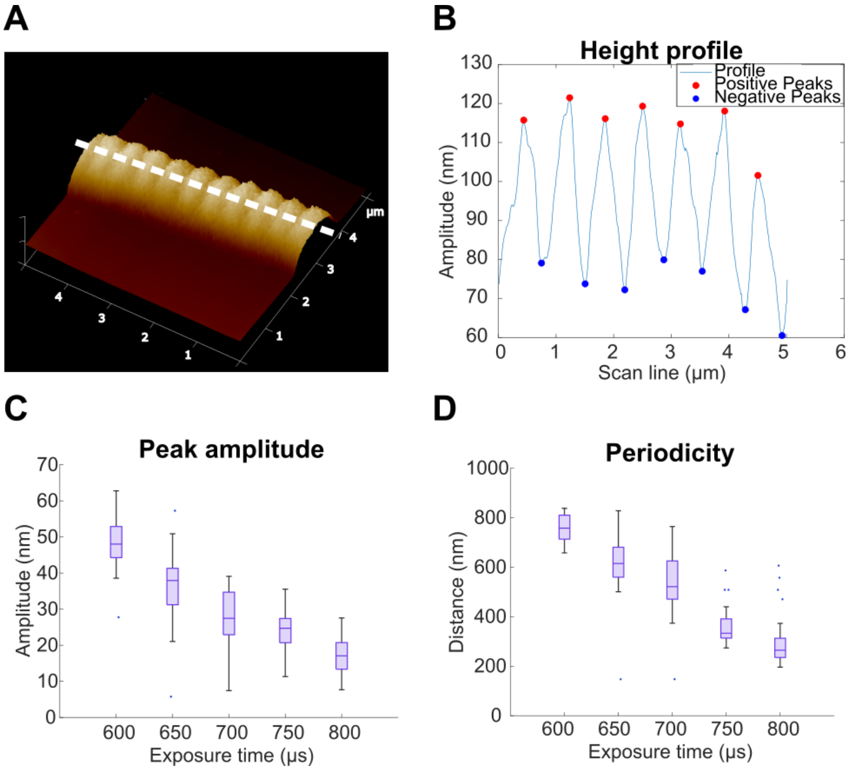

**Supplementary Figure 4 - Nanostructuration**

**A.** 3D view of a fiber and scan line (dashed white line). **B.** Height profile at the center of a fiber and positive and negative peak detection. **C.** Amplitude between successive positive and negative peaks for a range of exposure times. **D.** Nanostructuration periodicity (distance between successive positive peaks or successive negative peaks; N = 3 analyzed fibers for each condition). For all box plots, the central mark indicates the median, the edges of the box denote the 25th and 75th percentiles, the whiskers extend to the most extreme data points not considered outliers, and outliers are plotted individually (dots).

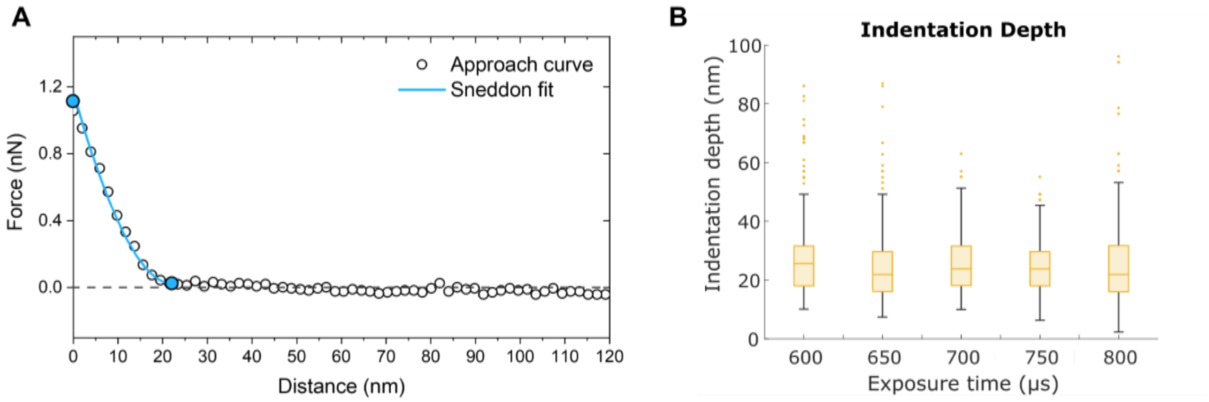

**Supplementary Figure 5 - Indentation depth and force-distance curve**

**A.** Typical force-distance curve obtained for the indentation experiment, here for $T_{exp}$ = 700 μs and an applied load of 1 nN. **B.** Contact point for an applied load of 1 nN. For all box plots, the central mark indicates the median, the edges of the box denote the 25th and 75th percentiles, the whiskers extend to the most extreme data points not considered outliers, and outliers are plotted individually (dots).

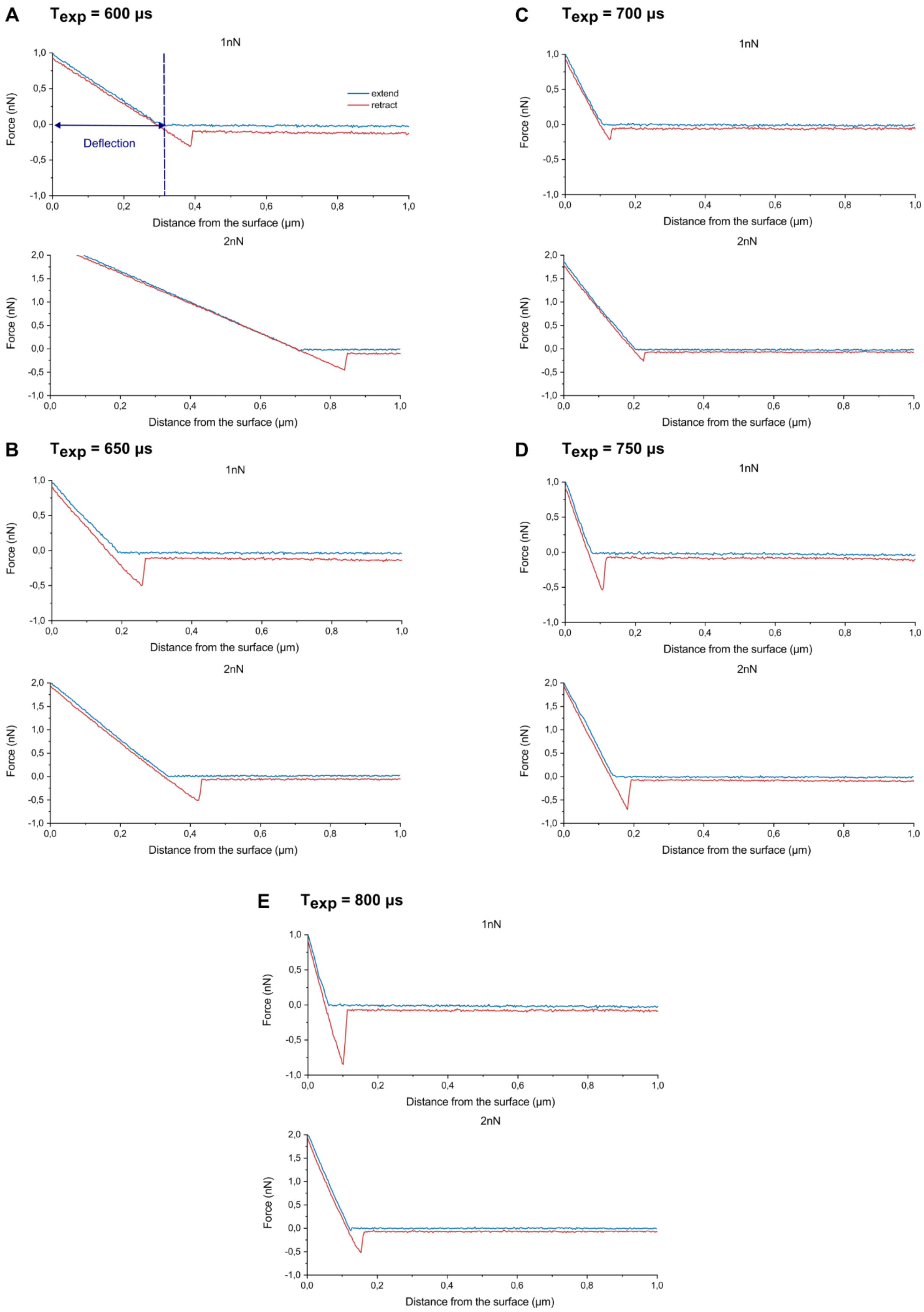
A
$T_{exp}$ = 600 µs
1nN
extend
retract
Deflection
Force (nN)
Distance from the surface (µm)
2nN
B
$T_{exp}$ = 650 µs
C
$T_{exp}$ = 700 µs
D
$T_{exp}$ = 750 µs
E
$T_{exp}$ = 800 µs

**Supplementary Figure 6 - Suspended fiber deflection experiment: force-distance curves**

Typical force-distance curves obtained when deflecting a fiber with a tipless AFM cantilever for **A.** $T_{exp}$ = 600 µs, **B.** $T_{exp}$ = 650 µs, **C.** $T_{exp}$ = 700 µs, **D.** $T_{exp}$ = 750 µs and **E.** $T_{exp}$ = 800 µs (applied load of top: 1 nN, bottom: 2 nN).

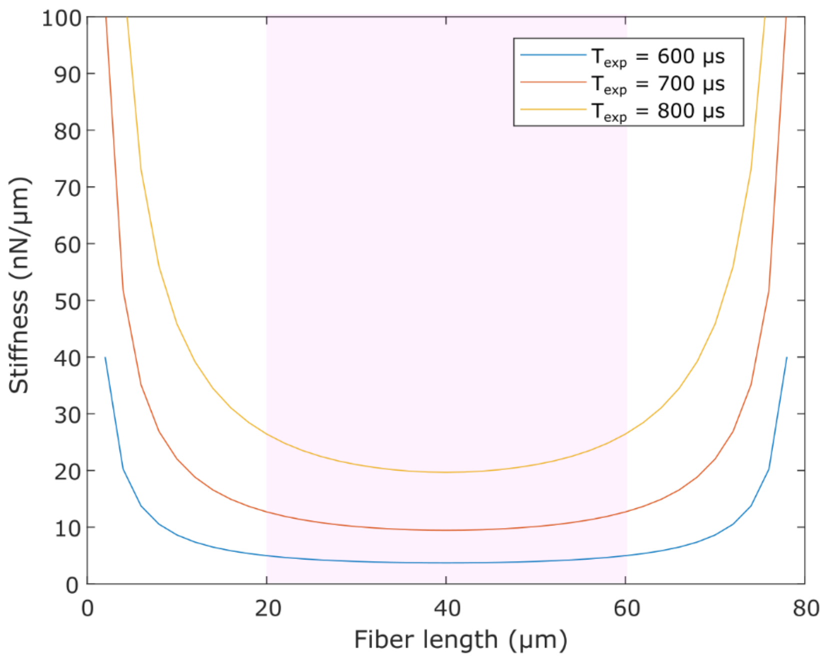

**Supplementary Figure 7 - Stiffness profile of a clamped beam under tension**

Theoretical stiffness profiles for photopolymerized fibers with $T_{exp}$ = 600 µs, 700 µs or 800 µs. The shaded area depicts the central portion of the fiber used for average force per node calculation, where the stiffness does not exceed a 30% increase compared to the stiffness at the midpoint of the fiber.

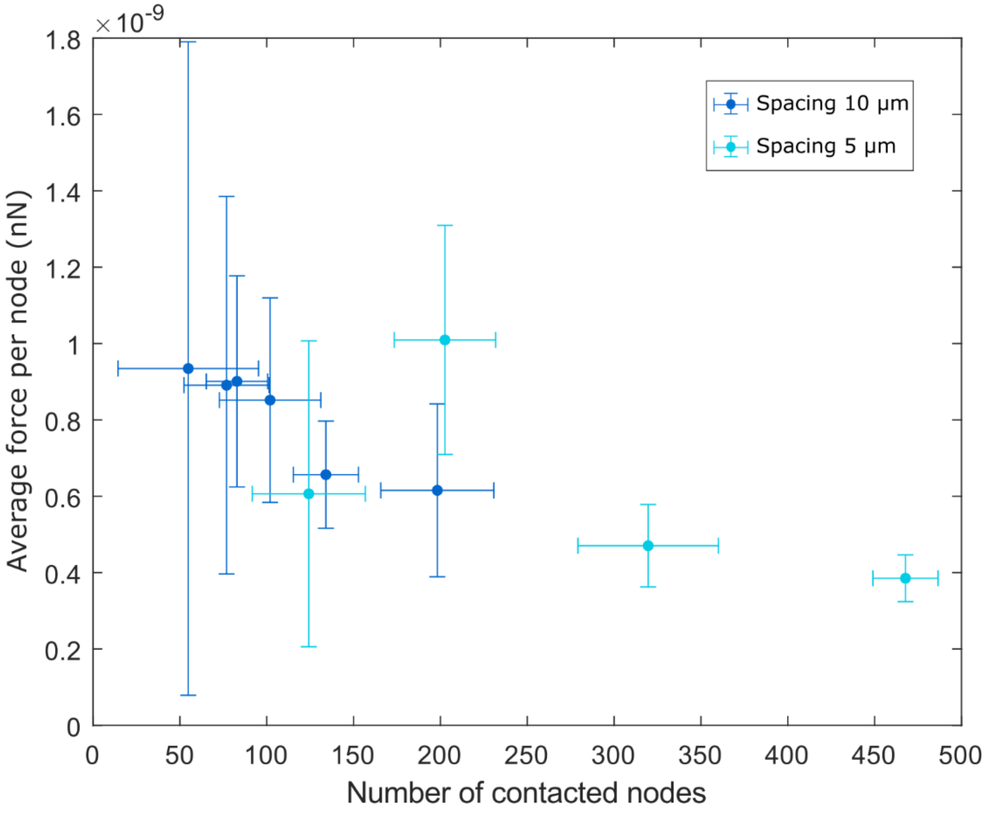

**Supplementary Figure 8 - Average force per node decreases with the number of contacted nodes**

Average force per node as a function of the number of contacted nodes for HUVECs on low-exposure fibers ($T_{exp}$ = 600 µs, k = 3.5 nN/µm) with lateral spacing of 5 µm and 10 µm. Each point represents the mean value over the whole timelapse for an individual cell and error bars correspond to the standard deviation.

### Supplementary Table

#### Supplementary Table 1 - Scanning speed

| **Exposure Time (µs)** | **Scanning speed (µm/s)** |
| --- | --- |
| 600 | 55.6 |
| 650 | 51.4 |
| 700 | 47.8 |
| 750 | 44.6 |
| 800 | 41.9 |

#### List of supplementary movies

**Supplementary movie S1:** Time lapse representing a NIH/3T3 Lifeact-GFP cell on a two-layer microscaffold (L = 80 µm) with highly deformable low-exposure fibers. The cell colormap codes for the *z*-position of the cell (purple = lower plane; blue = upper plane). Red: fibers. Total duration is 15 hours. Spinning-disk microscopy.

**Supplementary movie S2:** Time lapse representing a HUVEC Lifeact-GFP cell on a two-layer microscaffold (L = 80 µm) with highly deformable low-exposure fibers. The cell colormap codes for the z-position of the cell (purple = lower plane; blue = upper plane). Red: fibers. Total duration is 15 hours. Spinning-disk microscopy.

**Supplementary movie S3:** Time lapse representing HUVECs Lifeact-GFP on a two-layer large microscaffold (L = 200 µm) with highly deformable low-exposure fibers. The cell colormap codes for the *z*-position of the cell (purple = lower plane; blue = upper plane). Red: fibers. Total duration is 8 hours. Spinning-disk microscopy.

**Supplementary movie S4:** Time lapse representing HUVECs Lifeact-GFP on a two-layer large microscaffold (L = 200 µm) with moderately deformable high-exposure fibers. The cell colormap codes for the *z*-position of the cell (purple = lower plane; blue = upper plane). Red: fibers. Total duration is 8 hours. Spinning-disk microscopy.

**Supplementary movie S5:** Time lapse imaging of the development step following fiber polymerization during the microfabrication process. At t = 4 s, the monomer solution is washed with an excess of ethanol 100% and tension builds up within seconds. Total duration is 10 seconds.

**Supplementary movie S6:** 3D deflection map (left) and traction force map (right) of a NIH/3T3 cell on low-exposure fibers ($T_{exp}$ = 600 µs, k = 3.5 nN/µm, L = 80 µm, lateral spacing = 10 µm). For both maps, a maximum projection, top view and side view are represented. Bottom: evolution of contractility over time. Total duration is 15 hours.

**Supplementary movie S7:** 3D deflection map (left) and traction force map (right) of a NIH/3T3 cell on medium-exposure fibers ($T_{exp}$ = 700 µs, k = 8.3 nN/µm, L = 80 µm, lateral spacing = 10 µm). For both maps, a maximum projection, top view and side view are represented. Bottom: evolution of contractility over time. Total duration is 15 hours.

**Supplementary movie S8:** 3D deflection map (left) and traction force map (right) of a NIH/3T3 cell on high-exposure fibers ($T_{exp}$ = 800 µs, k = 17.6nN/µm, L = 80 µm, lateral spacing = 10 µm).

For both maps, a maximum projection, top view and side view are represented. Bottom: evolution of contractility over time. Total duration is 15 hours.

**Supplementary movie S9:** 3D deflection map (left) and traction force map (right) of a HUVEC cell on low-exposure fibers ($T_{exp}$ = 600 µs, k = 3.5 nN/µm, L = 80 µm, lateral spacing = 10 µm). For both maps, a maximum projection, top view and side view are represented. Bottom: evolution of contractility over time. Total duration is 17 hours.

**Supplementary movie S10:** 3D deflection map (left) and traction force map (right) of a HUVEC cell on low-exposure fibers ($T_{exp}$ = 600 µs, k = 3.5 nN/µm, L = 80 µm, lateral spacing = 5 µm). For both maps, a maximum projection, top view and side view are represented. Bottom: evolution of contractility over time. Total duration is 15 hours.

**Supplementary movie S11:** 3D deflection map (left) and traction force map (right) of a HUVEC cell on medium-exposure fibers ($T_{exp}$ = 700 µs, k = 8.3 nN/µm, L = 80 µm, lateral spacing = 10 µm). For both maps, a maximum projection, top view and side view are represented. Bottom: evolution of contractility over time. Total duration is 9 hours.

**Supplementary movie S12:** 3D deflection map (left) and traction force map (right) of a HUVEC cell on high-exposure fibers ($T_{exp}$ = 800 µs, k = 17.6 nN/µm, L = 80 µm, lateral spacing = 10 µm). For both maps, a maximum projection, top view and side view are represented. Bottom: evolution of contractility over time. Total duration is 17 hours.

**Supplementary movie S13:** 3D deflection map (left) and traction force map (right) of a macrophage on low-exposure fibers ($T_{exp}$ = 600 µs, k = 1.8 nN/µm, L = 160 µm, lateral spacing = 5 µm). For both maps, a maximum projection, top view and side view are represented. Bottom: evolution of contractility over time. Total duration is 130 minutes.

**Supplementary movie S14:** 3D deflection map (left) and traction force map (right) of a macrophage on low-exposure fibers ($T_{exp}$ = 600 µs, k = 1.8 nN/µm, L = 160 µm, lateral spacing = 10 µm). For both maps, a maximum projection, top view and side view are represented. Bottom: evolution of contractility over time. Total duration is 130 minutes.

**Supplementary movie S15:** 3D deflection map (left) and traction force map (right) of a macrophage on medium-exposure fibers ($T_{exp}$ = 700 µs, k = 4.1 nN/µm, L = 160 µm, lateral spacing = 5 µm). For both maps, a maximum projection, top view and side view are represented. Bottom: evolution of contractility over time. Total duration is 130 minutes.

**Supplementary movie S16:** 3D deflection map (left) and traction force map (right) of a macrophage on high-exposure fibers ($T_{exp}$ = 800 µs, k = 8.8 nN/µm, L = 160 µm, lateral spacing = 10 µm). For both maps, a maximum projection, top view and side view are represented. Bottom: evolution of contractility over time. Total duration is 130 minutes.

**Supplementary movie S17:** Time lapse representing a dendritic cell Lifeact-GFP cell on a two-layer microscaffold with low-exposure fibers ($T_{exp}$ = 600 µs, k = 3.5 nN/µm, L = 80 µm, lateral spacing = 10 µm). Pink: F-actin, green: fibers. Total duration is 20 minutes. Lattice Light-Sheet microscopy.

**Supplementary movie S18:** 3D deflection map (left) and traction force map (right) of a dendritic cell on low-exposure fibers ($T_{exp}$ = 600 µs, k = 3.5 nN/µm, L = 80 µm, lateral spacing = 10 µm), analysis of the Supplementary movie S17. For both maps, a maximum projection, top view and side view are represented. Bottom: evolution of contractility over time. Total duration is 20 minutes.
